# Infection- or vaccine mediated immunity reduces SARS-CoV-2 transmission, but increases competitiveness of Omicron in hamsters

**DOI:** 10.1101/2022.07.29.502072

**Authors:** Julia R. Port, Claude Kwe Yinda, Jade C. Riopelle, Zachary A. Weishampel, Taylor A. Saturday, Victoria A. Avanzato, Jonathan E. Schulz, Myndi G. Holbrook, Kent Barbian, Rose Perry-Gottschalk, Elaine Haddock, Craig Martens, Carl. I. Shaia, Teresa Lambe, Sarah C. Gilbert, Neeltje van Doremalen, Vincent J. Munster

**Author notes:** These first authors contributed equally (Julia R. Port, Claude Kwe Yinda). These senior authors contributed equally (Neeltje van Doremalen, Vincent J. Munster). New address: Chinese Academy of Medical Science Oxford Institute; Oxford Vaccine Group, Department of Paediatrics, University of Oxford, Oxford, UK.

## Abstract

Omicron has demonstrated a competitive advantage over Delta in vaccinated people. To understand this, we designed a transmission chain experiment using naïve, intranasally (IN) or intramuscularly (IM) vaccinated, and previously infected (PI) hamsters. Vaccination and previous infection protected animals from disease and virus replication after Delta and Omicron dual challenge. A gradient in transmission blockage was observed: IM vaccination displayed moderate transmission blockage potential over three airborne chains (approx. 70%), whereas, IN vaccination and PI blocked airborne transmission in >90%. In naïve hamsters, Delta completely outcompeted Omicron within and between hosts after dual infection in onward transmission. Although Delta also outcompeted Omicron in the vaccinated and PI transmission chains, an increase in Omicron competitiveness was observed in these groups. This correlated with the increase in the strength of the humoral response against Delta, with the strongest response seen in PI animals. These data highlight the continuous need to assess the emergence and spread of novel variants in populations with pre-existing immunity and address the additional evolutionary pressure this may exert on the virus.

## Introduction

In late 2019, SARS-CoV-2 spilled over into the human population, leading to the COVID-19 pandemic. Ongoing evolution in the human population resulted in the emergence of variants of concern (VOCs). Phenotypic changes that characterize VOCs are an increase in transmissibility, increase in virulence, change in clinical disease presentation, and/or decrease in effectiveness of public health and social measures or available diagnostics, vaccines, and therapeutics [1, 2]. Changes in the transmission phenotype can occur by a variety of adaptions including virus shedding dynamics, human behavior, host cell tropism, and entry. Furthermore, a large portion of the human population is no longer naïve to SARS-CoV-2 [3-5]. Immunity induced by previous exposure or vaccination have changed the susceptibility to infection and thus the evolutionary pressures on SARS-CoV-2. The emergence of VOCs is following almost a classic pattern in which the new VOC replaces the old VOC: this was observed for Alpha, Delta, and now Omicron. Whereas the initial replacements of previous VOCs by a new variant were due largely to an increase in the transmission potential of the virus, the transmission advantage of Omicron over Delta in humans is not fully understood [6]. Due to antigenic differences, the humoral response, especially the cross-reactivity of neutralizing antibodies from previous infections or vaccination against Omicron is poor [7-12]. Compared to Delta, Omicron is more likely to cause infections in a vaccinated population [13]. To better understand the directionality of SARS-CoV-2 evolution, it will be crucial to differentiate between the separate evolutionary pressures, including pre-existing immunity.

Previously, we have experimentally shown an increased aerosol transmission phenotype of SARS-CoV-2 Alpha over Lineage A [14, 15]. Here, we are using infection- or vaccine-mediated immunity to model the impact of this evolutionary pressure on the transmission of the Delta and Omicron VOCs.

## Results

### Decreased spike-mediated entry and delayed shedding kinetics of Omicron over Delta

Delta and Omicron VOCs have several observed mutations in the spike (S) protein, including the receptor binding domain (RBD), N-terminal domain (NTD), and the S1/S2 cleavage site (**Figure 1 A**). To determine if these changes in S might affect the behavior of these variants in the hamster model, we modeled changes on the structure of the RBD – ACE2 complex and evaluated entry of the variants using our VSV-pseudotype entry assay. We previously observed that of the residues on ACE2 that directly participate in RBD binding [16], two contact residues differ between human and hamster ACE2 [14]. In hamster ACE2, the histidine (H) and methionine (M) at position 34 and 82, respectively, are replaced by glutamine (Q) and asparagine (N) (**Figure 1 B**, red). Two of the mutations in Omicron, K417N and Q493R, are in close proximity to the H34Q substitution observed in hamster ACE2 (**Figure 1 B**) and could potentially lead to altered interactions with ACE2 at this location.

**Figure 1.**
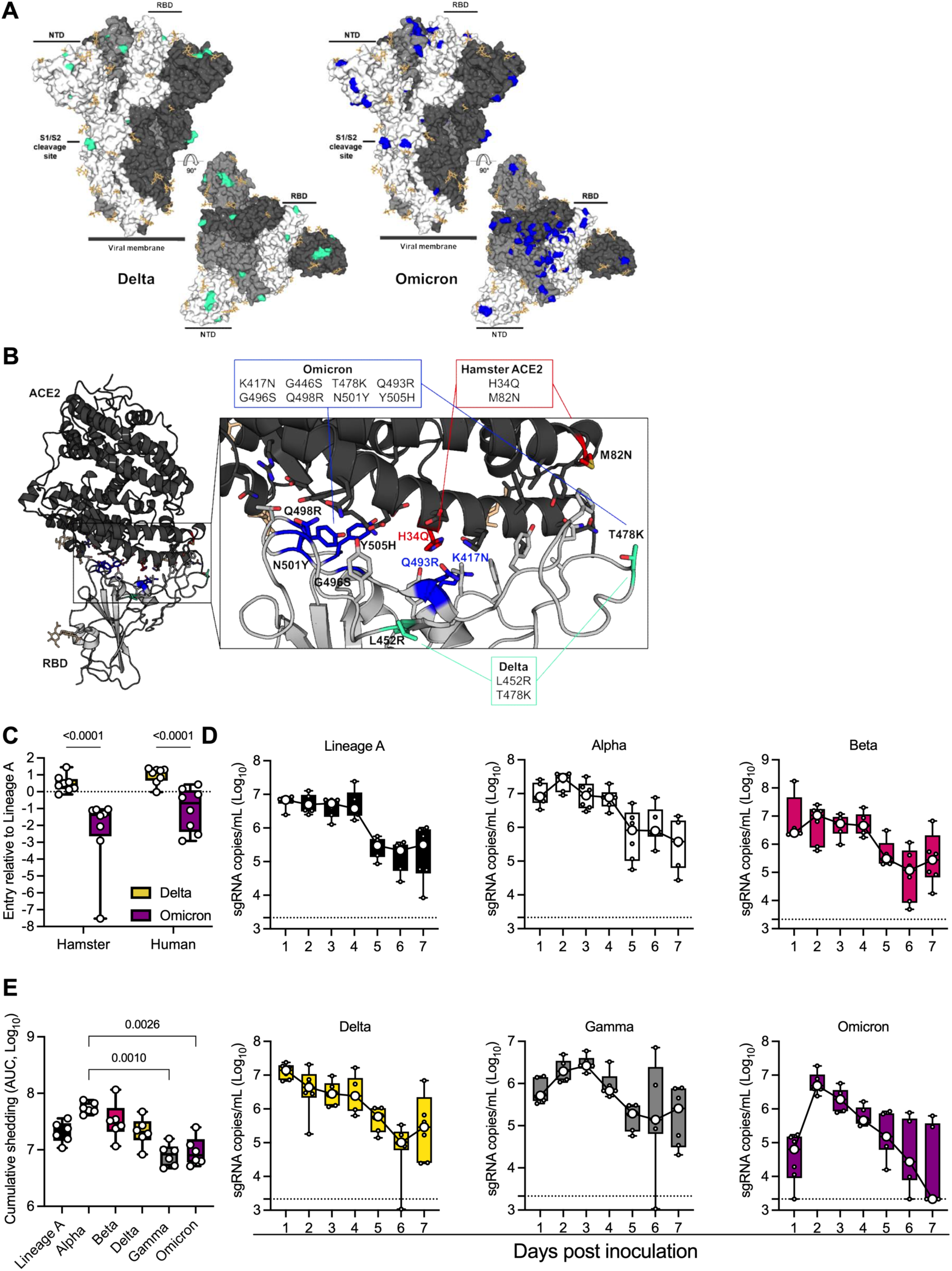
Comparison of SARS-CoV-2 variants Omicron and Delta infection in the Syrian hamster. **A**. Mutations observed in the SARS-CoV-2 Delta and Omicron VOCs are highlighted on the structure of SARS-CoV-2 spike (compared to Lineage A, PDB 6ZGE, [52]). The spike trimer is depicted by surface representation with each protomer colored a different shade of gray. The residues at the positions of the spike protein mutations observed in the Delta and Omicron SARS-CoV-2 VOCs are colored teal green (Delta) and blue (Omicron). The receptor binding domain (RBD), N-terminal domain (NTD), and cleavage site are annotated. N-linked glycans are shown as light, orange-colored sticks. **B**. The structure of the Alpha VOC RBD and human ACE2 complex (PDB 7EKF, [53]) is depicted with cartoon representation. ACE2 is colored dark gray and the RBD is colored light gray. N-linked glycans are shown as light, orange-colored sticks. A box reveals a close-up view of the RBD-ACE2 binding interface. Side chains of the residues participating in the interaction, as identified and described by Lan, et al [16] are shown as sticks. The residues within the RBD that are mutated in the Delta and Omicron VOCs are colored teal green (Delta) and blue (Omicron). Residue T478 is mutated in both Delta and Omicron VOCs but is colored teal green in the figure. Though they do not participate directly in the ACE2 interface, the sidechains of residues L452 and T478 are also shown. The residues that differ between human and hamster ACE2 within the interface are colored red. **C**. BHK cells expressing either human ACE2 or hamster ACE2 were infected with pseudotyped VSV reporter particles with the spike proteins of Delta or Omicron. Relative entry to a lineage A control is depicted. Whisker-plots depicting median, min and max values, and individual values, N = 8, ordinary two-way ANOVA, followed by Šídák’s multiple comparisons test. **D**. Viral load as measured by sgRNA in oropharyngeal swabs collected at 1-7 days post intranasal 1,000 TCID_50_ inoculation with Lineage A, Alpha, Beta, Delta, Gamma or Omicron. Whisker-plots depicting median, min and max values, and individual values, N = 6 (3 males and 3 females). **E**. Cumulative sgRNA shedding for each variant. Area under the curve for data shown in D. Kruskal-Wallis test. P-values stated were significant (<0.05).

Next, we evaluated cellular entry by the S protein of the Delta and Omicron VOCs compared to the ancestral Lineage A S protein using the VSV-pseudotype entry assay system in baby hamster kidney (BHK) cells expressing either human or hamster ACE2 (**Figure 1 C**). For human ACE2, the entry of the Omicron S was similar to that of the ancestral Lineage A S but significantly lower than that of the Delta variant (mean difference = 1.397-fold entry over Lineage A, p <0.0001, N = 8, two-way ANOVA, followed by Šídák’s multiple comparisons test). For hamster ACE2, we observed a 1.55 mean difference between Delta and Omicron (p <0.0001, N = 8, two-way ANOVA, followed by Šídák’s multiple comparisons test).

Based on the *in silico* and *in vitro* data, we evaluated whether the displayed phenotype of Omicron would result in a change in respiratory shedding in the Syrian hamster model in comparison to Alpha, Beta, Gamma, Delta, or Lineage A (**Figure 1 D**). Six hamsters per group were inoculated with 10^3^ TCID_50_ of SARS-CoV-2 variants via the intranasal (IN) route. Oropharyngeal swabs were taken for 7 days. Sub-genomic (sg)RNA shedding peaked on day 1 post-inoculation for Lineage A and Delta in contrast to Alpha, Beta, and Omicron, for which median peak shedding was highest on day 2. Only Gamma showed peak shedding on day 3 post inoculation. Median peak shedding for all variants ranged between 10^6^ and 10^7^ sgRNA copies/mL. When comparing the cumulative shedding (area under the curve (AUC)), animals inoculated with Alpha shed significantly more than those inoculated with Gamma and Omicron variants (**Figure 1 E**, N = 6, Kruskal-Wallis test, followed by Dunn’s multiple comparisons test, p = 0.0010 and 0.0026, respectively).

### Contact and airborne transmission in naïve Syrian hamsters

In the human population, Omicron replaced Delta as the most prevalent variant [17]. To understand whether this is due to an increase in transmissibility, we compared transmission of Delta and Omicron in transmission chains in naïve hamsters. We performed contact and airborne transmission chain experiments over two or three generations (1:1 ratio between donors and sentinels) and repeated these chains three times (**Figure 2 A**). Donors were intranasally inoculated with a 1:1 mixture of Delta and Omicron. One day later, generation 1 sentinels (sentinels 1) were exposed to the donors for 48h, followed by exposure of sentinels 2 to sentinels 1 for 48h, and finally exposure of sentinels 3 to sentinels 2 for 72h. Each exposure was started on 2 DPI/DPE relative to the previous chain. Oropharyngeal swabs were collected from all animals at 2, 3, and 5 DPI/DPE, and lung and nasal turbinate samples were harvested at 5 DPI/DPE.

**Figure 2.**
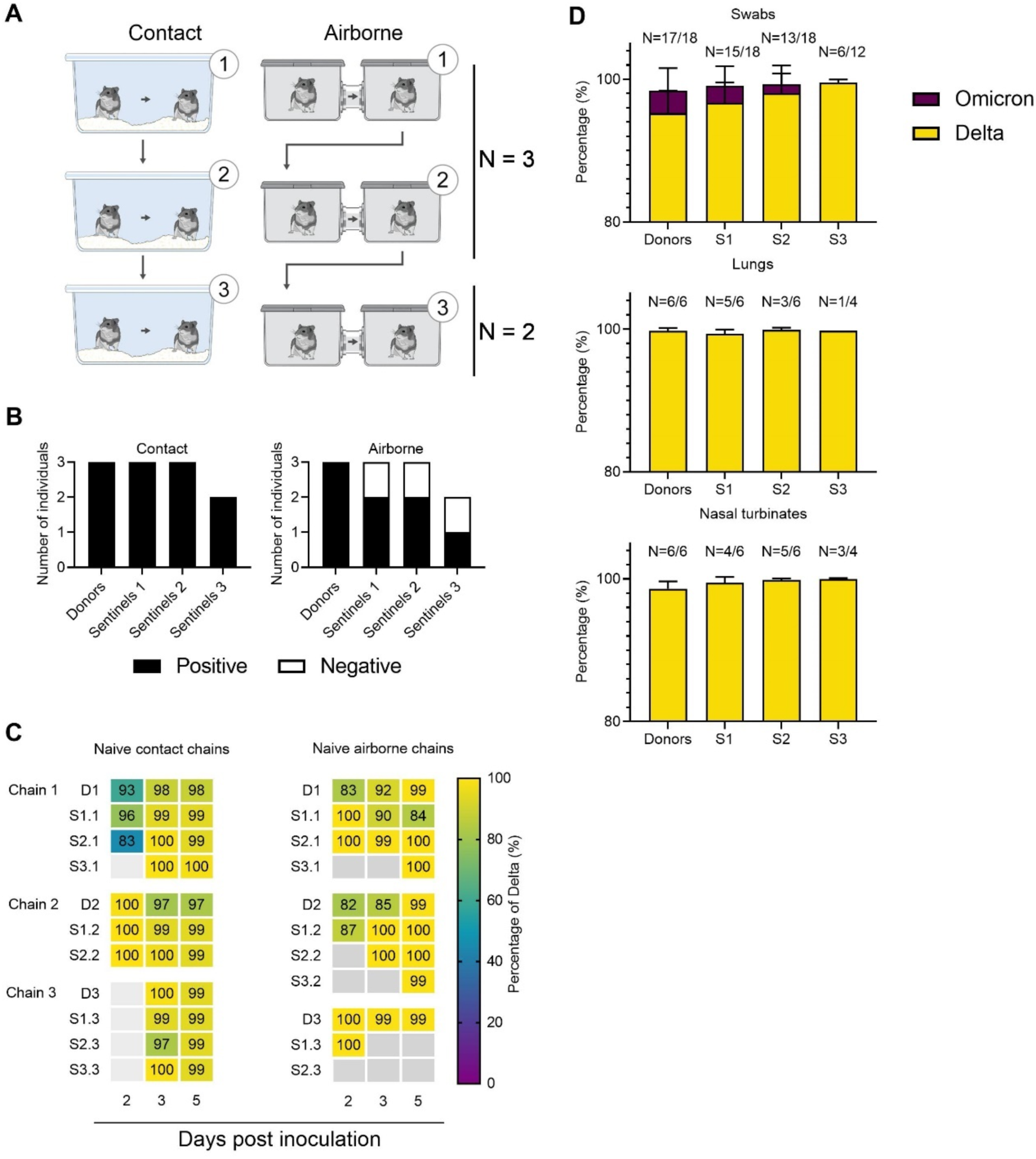
Transmission competitiveness of Delta and Omicron in a naïve hamster population. Chain transmission in naïve Syrian hamsters assessing the competitiveness of Delta and Omicron over three transmission events. **A**. Donor animals (N = 6) were inoculated with a total of 10^4^ TCID_50_ of Delta and Omicron (1:1 ratio) via the IN route, and three groups of sentinels (sentinels 1 (N = 6), 2 (N = 6) and 3 (N = 4)) were subsequently exposed. Half were exposed by direct contact (housed in the same cage), and half at 16.5 cm distance (airborne exposure). Animals were exposed at a 1:1 ratio; exposure occurred 24h post inoculation (Donors ⟶ sentinels 1) and 48h post exposure for subsequent groups (sentinels ⟶ sentinels). **B**. Summary of infection status for the donors and sentinels. Oropharyngeal swabs were taken on 2, 3, and 5 DPI/DPE, and lungs and nasal turbinates were collected at day 5 DPI/DPE. Individuals were considered infected, if 2 out of 5 samples were positive for sgRNA (oral swab, lung or nasal turbinate). Bar charts depict summary of individuals, divided into contact and airborne chains. **C**. The receptor binding domain of the SARS-CoV-2 spike was sequenced for all sgRNA positive swabs collected at 2, 3, and 5 DPI/DPE. Heatmap representing all sgRNA positive swab samples from each individual for each chain and showing the percentage of Delta detected. Colors refer to legend on right (D = donor, S = sentinel), grey = no sgRNA present in the sample or sequencing unsuccessful. **D**. Overall percentage of Delta and Omicron in all sgRNA positive samples in each group, separated by sample type. Bar charts depicting median and 95% CI. Number of sgRNA positive samples over all samples analyzed is indicated on top. Yellow = Delta, purple = Omicron.

Animals were considered infected, when 2 out of 5 samples collected had detectable sgRNA, a marker of viral replication. In the naïve direct contact chains, all animals became infected (**Figure 2 B, left panel**). In contrast, 2 out of 3 of the sentinels 1 and sentinels 2 hamsters, and 1 out of 2 sentinels 3 hamsters became infected in the airborne chains. When excluding sgRNA negative samples, no significant difference was observed in the median viral sgRNA titers in lungs, nasal turbinates or swabs on day 2 or 3 between donors and sentinels with both routes of transmission combined **(Supplementary Figure 1 and 2, Supplementary Table 2)**.

We analyzed the relative composition of each of the VOCs in all sgRNA positive samples by NGS. Delta outcompeted Omicron both within and between hosts (**Figure 2 C**). Across all sgRNA positive swabs in donors and sentinels, Delta comprised >98% of viral sequences, though some individual variation was observed in swabs. The percentage of Delta increased with each subsequent transmission chain in swabs (median percentage Delta in donors = 98% (99.9 – 81.8 95% CI); sentinels 1 = 99 (100 – 84.4 95% CI); sentinels 2 = 99.5% (100 – 83 95% CI); and sentinels 3 = 99.8 (99.9 – 99 95% CI). No Omicron was detected in lungs or nasal turbinates (**Figure 2 D**).

### Previous exposure or vaccine-induced pre-existing immunity against Lineage A or Delta reduces virus replication, shedding and lung pathology after reinfection

Next, we compared the impact of pre-existing immunity on the contact and airborne competitiveness of Delta and Omicron (**Figure 3 A**). Pre-existing immunity was achieved by IN or intramuscular (IM) vaccination with AZD1222, or previous infection with Delta. 16 hamsters per group were immunized with AZD1222 (ChAdOx1 nCoV-19, 2.5 × 10^8^ IU/animal) or exposed via direct contact to IN-inoculated animals one day after inoculation (5:1 sentinel : donor ratio, previously infected group (PI)). In all vaccinated and PI animals, seroconversion was confirmed after 21 days (**Figure 4 A**). 28 days after immunization via vaccination or infection, 6 animals per group were challenged via the IN route using 10^4^ TCID_50_ SARS-CoV-2 (1:1 mixture, Delta and Omicron variants).

**Figure 3.**
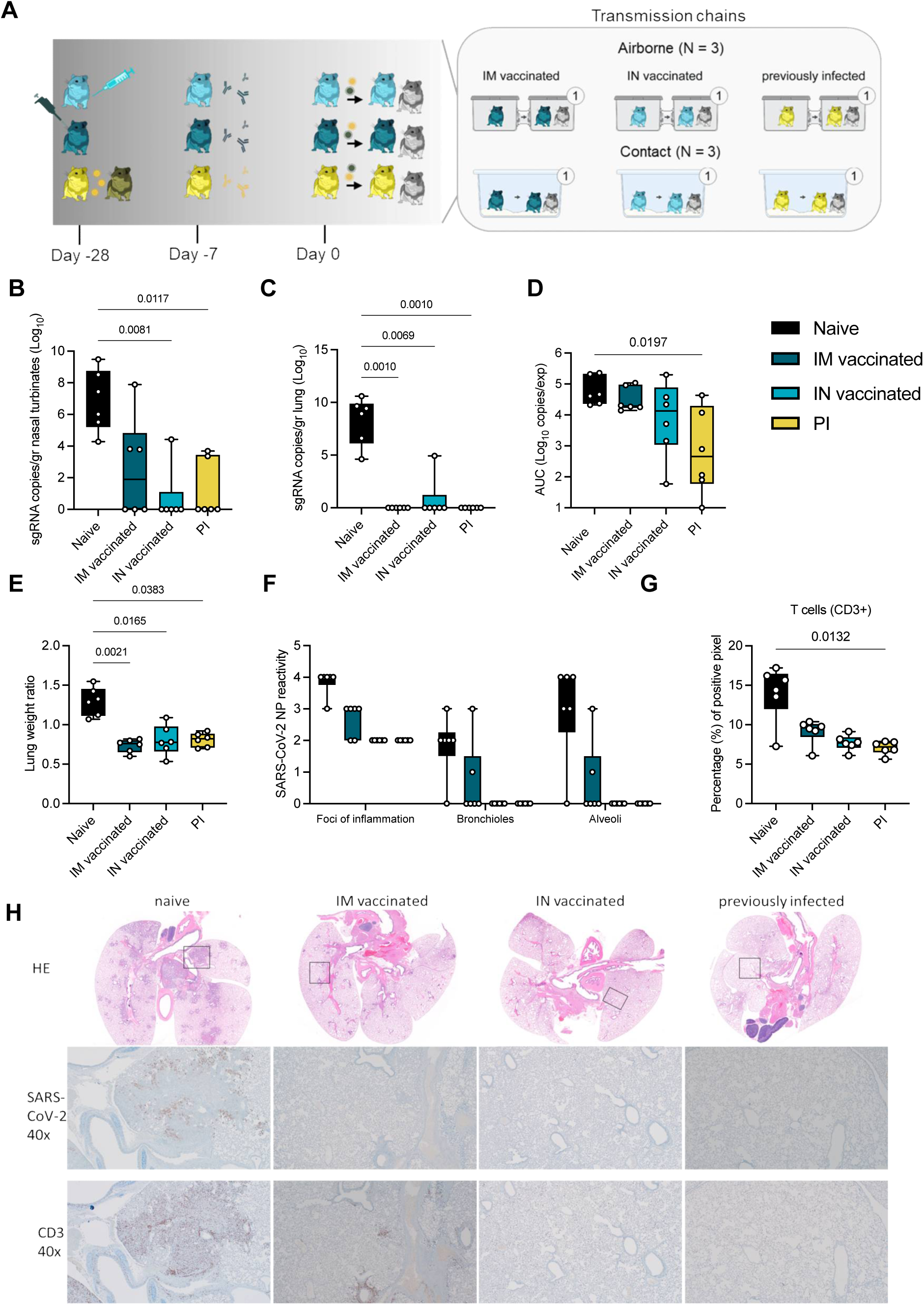
Reduction of disease severity and shedding through pre-existing immunity. **A**. Schematic. Hamsters were either vaccinated IN or IM against lineage A or experienced a previous infection with Delta through contact exposure to IN inoculated hamsters. Immune status was confirmed after 21 days. Transmission competitiveness in these populations was investigated at least 28 days post vaccination or infection. Donor animals (N = 6 for each group) were inoculated with a total of 10^4^ TCID_50_ of Delta and Omicron via the IN route (1:1 ratio), and sentinels 1 (N = 6) were exposed 24h later. For each transmission event, a naïve control animal was also exposed. Half were exposed by direct contact (housed in the same cage), and half at 16.5 cm distance (airborne exposure). Animals were exposed at a 1:1:1 ratio, and exposure occurred on day 1 and lasted for 48 hours. **B.C**. Tissue samples were collected at 5 DPI/DPE for donors. Donor sgRNA in lungs and nasal turbinates. Whisker-plots depicting median, min and max values, and individual values, N = 6, ordinary two-way ANOVA, followed by Šídák’s multiple comparisons test. **D**. Cumulative shedding. Area under the curve (AUC) of sgRNA measured in oral swabs taken on 2,3, and 5 DPI. Whisker-plots depicting median, min and max values, and individual values, N = 6, ordinary two-way ANOVA, followed by Šídák’s multiple comparisons test. **E**. Lung weights (lung:body weight ratio). **F**. SARS-CoV-2 reactivity measured by immunohistochemistry targeting SARS-CoV-2 nucleoprotein (NP) in upper and lower respiratory tract. Whisker-plots depicting median, min and max values, and individual values, N = 6, ordinary two-way ANOVA, followed by Šídák’s multiple comparisons test. **G**. T-cell infiltration into the lung, measure by CD3 antigen presence and positive pixel quantification. Whisker-plots depicting median, min and max values, and individual values, N = 6, Kruskal-Wallis test. black = naïve, dark blue = IM vaccinated, light blue = IN vaccinated, yellow = PI. P-values stated were significant (<0.05). **H**. Lung pathology. Top = HE stains, middle = IHC for nucleoprotein, bottom = IHC for CD3. Squares indicate area of magnification.

**Figure 4.**
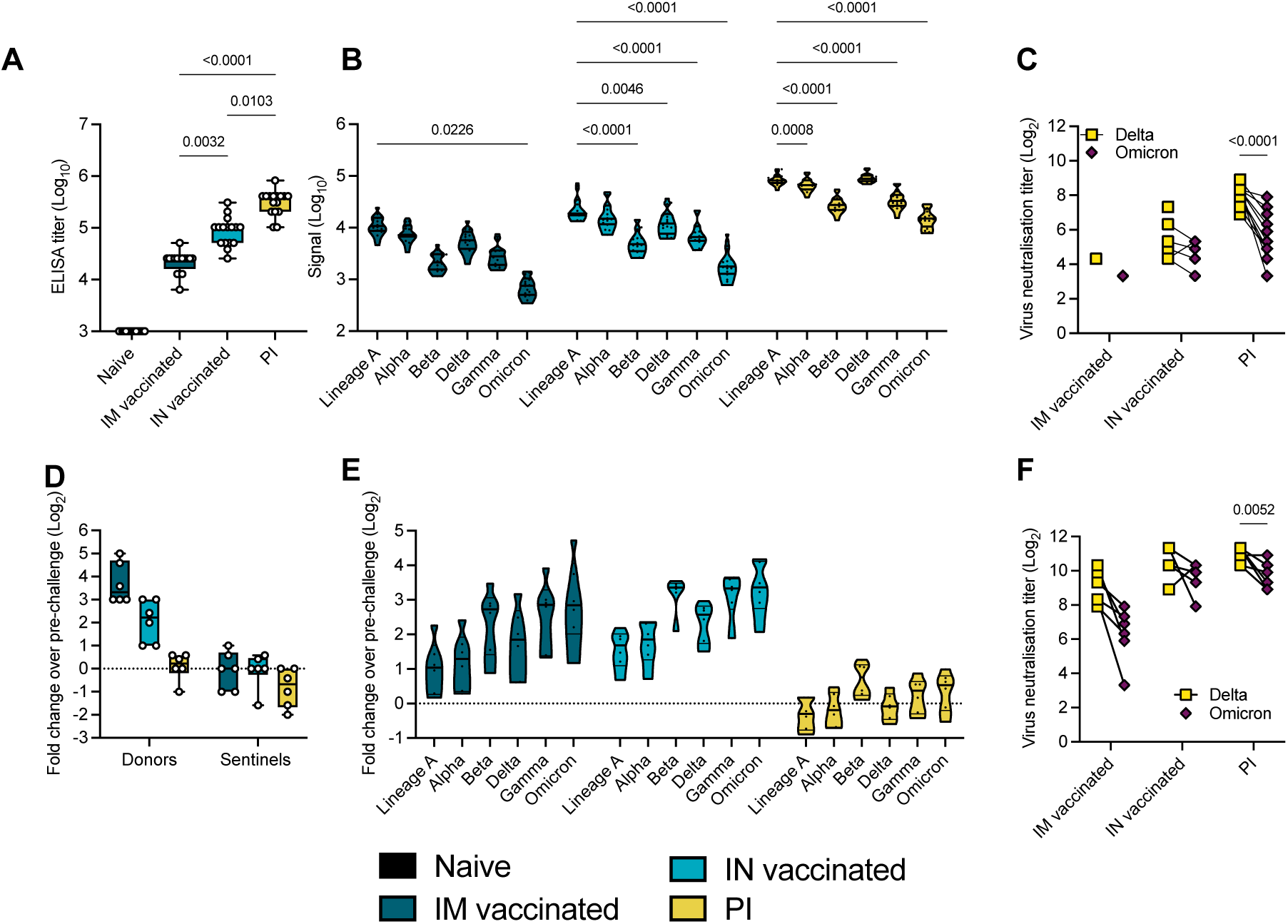
Variant specific infection- or vaccine mediated humoral immunity. Serology in IN or IM vaccinated and PI hamsters pre- and post-challenge with Delta/Omicron. Serum was collected 21 days post vaccination or infection with Delta, and on 5 DPI/DPE. **A**. Anti-spike IgG response, measured by ELISA. Whisker-plots depicting median, min and max values, and individuals. Kruskal-Wallis test, N = 16. **B**. Cross-reactivity of the IgG response, measured by Meso QuickPlex. Violin plots depicting median, quantiles, and individual values. Two-way ANOVA, followed by Šídák’s multiple comparisons test. N = 16. **C**. Individual neutralizing antibody titers against Delta and Omicron. Points connected by lines indicate the same animal. Two-way ANOVA, followed by Šídák’s multiple comparisons test. N = 16. **D**. Change in overall anti-spike IgG response after challenge (donors and sentinels). Whisker-plots depicting median, min and max values, and individual values. Change in titer is represented as Log_2_ (fold change over pre-challenge value). Dotted line indicates no change in titer. Kruskal-Wallis test, N = 6. **E**. Change in cross-reactivity after challenge/re-infection in donors. Violin plots depicting median, quantiles, and individual values. Change in titer is represented as Log_2_ (fold change over pre-challenge value). Dotted line indicates no change in titer. Two-way ANOVA, followed by Šídák’s multiple comparisons test. N = 16. **F**. Individual neutralizing antibody titers of donors against Delta and Omicron after challenge. Points connected by lines indicate the same animal. Two-way ANOVA, followed by Šídák’s multiple comparisons test. N = 6. black = naïve, dark blue = IM vaccinated, light blue = IN vaccinated, yellow = previously infected. P-values stated were significant (<0.05).

First, we assessed the impact of pre-existing immunity on viral replication and pathogenicity in the naïve, IN, IM, or PI donors. In naïve animals, virus replication was observed in nasal turbinates (median = 6.873 sgRNA copies/gr (Log_10_)) and lung tissue (median = 8.303 sgRNA copies/gr (Log_10_)). In contrast, viral RNA load was significantly reduced or absent in IM, IN, and PI groups as compared to naïve donors (Kruskal-Wallis test, followed by Dunn’s multiple comparison test, N = 6; lung: p = 0.0010, 0.0069, 0.0010, respectively; nasal turbinates: p = 0.1479, 0.0081, 0.0117, respectively). In lung tissue, sgRNA was only detected in 1 out of 6 animals in the IN group (4.93 sgRNA copies/gr (Log_10_)), but not in the other groups. sgRNA was detected in 3 out of 6 nasal turbinate samples in the IM donors (median = 2.581 sgRNA copies/gr (Log_10_)), 1 out of 6 in the IN group (4.423 sgRNA copies/gr (Log_10_)), and 2 out of 6 in the PI group (median = 1.173 sgRNA copies/gr (Log_10_), **Figure 3 B/C**).

Vaccination and previous infection reduced overall respiratory shedding. We measured sgRNA on 2, 3, and 5 DPI in oral swabs. Cumulative virus burden (area under the curve (AUC)) in oral swabs was marginally reduced after IM vaccination (median AUC (Log_10_) = 19,489, p = 0.999, N = 6, Kruskal-Wallis test, followed by Dunn’s multiple comparison test), moderately reduced after IN vaccination (median AUC (Log_10_) = 13,470, p = 0.4347), and significantly reduced in the PI group (median AUC (Log_10_) = 454.4, p = 0.0197), compared to naïve animals (median AUC (Log_10_) = 43,618) (**Figure 3 D**).

We compared the severity of lung disease as measured by the lung:body weight ratio (**Figure 3 E**). In the donor hamsters, previously established immunity reduced the lung:body weight ratio significantly after challenge (naïve = 1.296, IM = 0.7343, IN = 0.8030, PI = 0.8077, N = 6, Kruskal-Wallis test, followed by Dunn’s multiple comparison test, run against the naïve group, p = 0.021, p = 0.0165, and p = 0.0383, respectably). Hamsters from the naïve group developed lesions typical of SARS-CoV-2 in this model [18] (**Figure 3 H, Supplementary Table 1**). SARS-CoV-2 nucleoprotein immunoreactivity, a measurement of viral presence, ranged from moderate to numerous in both bronchi and alveoli and was especially apparent at the periphery of foci of pneumonia (**Figure 3 F and H**). CD3 immunoreactivity, a measurement of T-cell infiltration, was greatly increased in foci of inflammation and pneumonia in the lung (**Figure 3 G**). IM vaccination decreased the disease severity, as previously described [19, 20], which was accompanied by decreased antigen presence and T-cell infiltration compared to naïve animals. The majority of CD3 immunoreactive T-cells were located adjacent to bronchioles and blood vessels. In contrast, pathology in the IN vaccinated and PI hamsters was negligible and limited to scant inflammation and terminal airway reactivity, with no detectable virus presence and consistently lower T-cell numbers than the naïve animals. Surprisingly, no difference in B-cell infiltration was observed, as measured by PAX5 staining between the groups (**Supplemental Table 1**).

### Pre-existing humoral immunity against lineage A or Delta offers minimal neutralizing cross-reactivity against Omicron

To quantify the immune pressure against Omicron in our groups, IgG anti-spike responses were analyzed. All animals seroconverted by day 21 post vaccination or infection with Delta (**Figure 4 A**). Compared to IM vaccination, IN vaccination led to 4-fold higher humoral responses (median titer IM vaccinated = 25,600; median titer IN vaccinated = 102,400, p = 0.0032, Kruskal-Wallis test, followed by Dunn’s multiple comparison test, N = 16). PI hamsters had significantly higher titers (median = 409,600) than both IN vaccinated (p = 0.0103) and IM vaccinated (p < 0.0001) hamsters. To better assess the production of binding antibodies in the IM, IN, and PI groups, we analyzed the positive sera on a MESO QuickPlex panel [18] (**Figure 4 B**). In the IM and IN groups, the highest median signal was seen with an antibody response to Lineage A (IM group = 10788.75; IN group = 18692.00; PI group = 81855.75), which supports results from previous studies with vaccines against Lineage A [21]. While the response was strongest against Delta in the PI group, the overall response pattern to different variants was similar across all three groups. The median response signal against Omicron was consistently lower than against Delta. Next, a live virus neutralization assay was performed with the Delta and Omicron VOCs. Consistent with the ELISA and Meso QuickPlex results, neutralizing antibody titers were highest in the PI group, which neutralized Delta >10-fold better than Omicron (p < 0.0001, N = 6, two-way ANOVA followed by Šídák’s multiple comparisons test) (**Figure 4 C**). In the IN vaccinated hamsters, 9 out of 16 animals showed no neutralizing antibodies against the Omicron variant. Of IM vaccinated hamsters, 14 out of 16 had no neutralization of the Delta variant and 15 out of 16 had no neutralization of the Omicron variant. These results indicate that prior infection produced the most robust neutralizing antibody response, and that this response is more effective against the Delta variant than the Omicron variant.

Next, for each donor hamster, the fold change in post-challenge antibody titer relative to their pre-challenge baseline was calculated in samples collected at 5 DPI (**Figure 4 D**). IM donors experienced a median 10-fold change which was higher as compared to IN donors, which had a median fold-change value of 4.7, and PI donors, which had a titer fold-change of 1.2. These results indicate greater increases in IgG titers in response to challenge in hamsters that had lower antibody titers at baseline. Variant specific fold-change increase confirmed this finding. Interestingly, the challenge with the 1:1 Omicron/Delta inoculum induced the same affinity maturation profile across groups. The largest fold-change increase was observed for the antigenically most distant variants as compared to the initial priming variant, namely Beta, Gamma, and Omicron (**Figure 4 E**). A live virus neutralization assay was performed against the Omicron and Delta variants. Intriguingly, the relative difference in fold-change neutralization capacity against Omicron compared to Delta decreased. Yet, all groups maintained higher levels of neutralizing antibodies against Delta than Omicron, with median titers against Delta four-fold higher than Omicron in PI animals (p = 0.0052, N = 6, two-way ANOVA followed by Šídák’s multiple comparisons test) (**Figure 4 F**).

### Pre-existing humoral immunity protects against contact and airborne transmission

We hypothesized that under pre-existing immune pressure, the competition between Delta and Omicron would favor Omicron due to the larger antigenic distance relative to the previous lineages of SARS-CoV-2. To test this, groups of animals with vaccine- or infection-induced pre-existing immunity were used in a contact and airborne transmission experiment. Twenty-four hours after SARS-CoV-2 (1:1 mixture, Delta and Omicron variant) challenge, donors were co-housed with one naïve sentinel and one immunized sentinel (sentinels 1, 1:1:1 ratio) for 48 hours. This enabled us to compare airborne and contact transmission between donors and sentinels (N = 3, 1:1 ratio) for IM vaccinated, IN vaccinated, and PI hamsters (**Figure 4 A, Supplementary Figure 1 and 2, Supplementary Table 2**). An animal was considered infected if 2 out of 5 samples (either a swab, nasal turbinates, or lung tissue sample) had detectable sgRNA. In donor animals which were directly inoculated with 10^4^ TCID_50_ of virus, IN vaccination and PI reduced virus replication compared to IM vaccination. All IM vaccinated donors became infected. In contrast, 5 out of 6 donors in the IN vaccinated group, and 3 out of 6 donors in the PI group became infected.

We then assessed infection in the naïve and immunized sentinels 1 after contact transmission. For IM vaccination, 2 out of 3 naïve sentinels and 2 out of 3 immunized sentinel 1 hamsters became infected. Contact transmission was further reduced in the IN vaccinated and PI groups. For both chains, 2 out of 3 donors were infected, but only 1 out of 3 immunized sentinels 1 and no naïve sentinels 1 were infected.

Reduction in transmission was more prominent in the airborne chains. Only 1 out of 3 immunized sentinels 1 was infected in the IM airborne chains, while 2 out of 3 naïve sentinel hamsters became infected. In the IN vaccinated airborne chains, 1 out of 3 immunized sentinels 1 and no naïve sentinel 1 hamsters were infected. In the PI chains, no immunized or naïve sentinel 1 became infected (**Figure 5 A)**. Due to the importance of airborne transmission, and the increased reduction in airborne transmission already observed between donors and sentinels, we decided to take two airborne chains out to sentinels 3 (as described above for the naïve hamsters: donors ⟶ sentinels 1 ⟶ sentinels 2 ⟶ sentinels 3). In the IM vaccinated group, 1 out of 2 immunized sentinel 2 animals, but no naïve sentinel 2, no immunized sentinel 3, and no naïve sentinel 3 were infected. In the IN vaccinated group, no sentinel 2 and no sentinel 3 became infected. In the PI group, 1 out of 2 immunized sentinel 2 animals, but no naïve sentinel 2, nor any sentinel 3 became infected. We compared the airborne transmission efficiency between naïve, IM vaccinated, IN vaccinated, and PI hamsters. using the data across all transmission events, including immunized and naïve sentinels. For naïve hamsters, the airborne transmission efficiency was 63% (percentage of all transmission events resulting in an infected sentinel/all transmission events). Both vaccination and previous infection reduced this efficacy. While IM vaccination reduced of airborne transmission to 29% (p = 1.870, Fisher’s exact test, two sided: Odds ratio = 4.167), both IN vaccination (p = 0.0109, Fisher’s exact test, two sided: Odds ratio = 21.67) and PI (p = 0.0109, Fisher’s exact test, two sided: Odds ratio = 21.67) reduced it to 7% (**Figure 5 B**).

**Figure 5.**
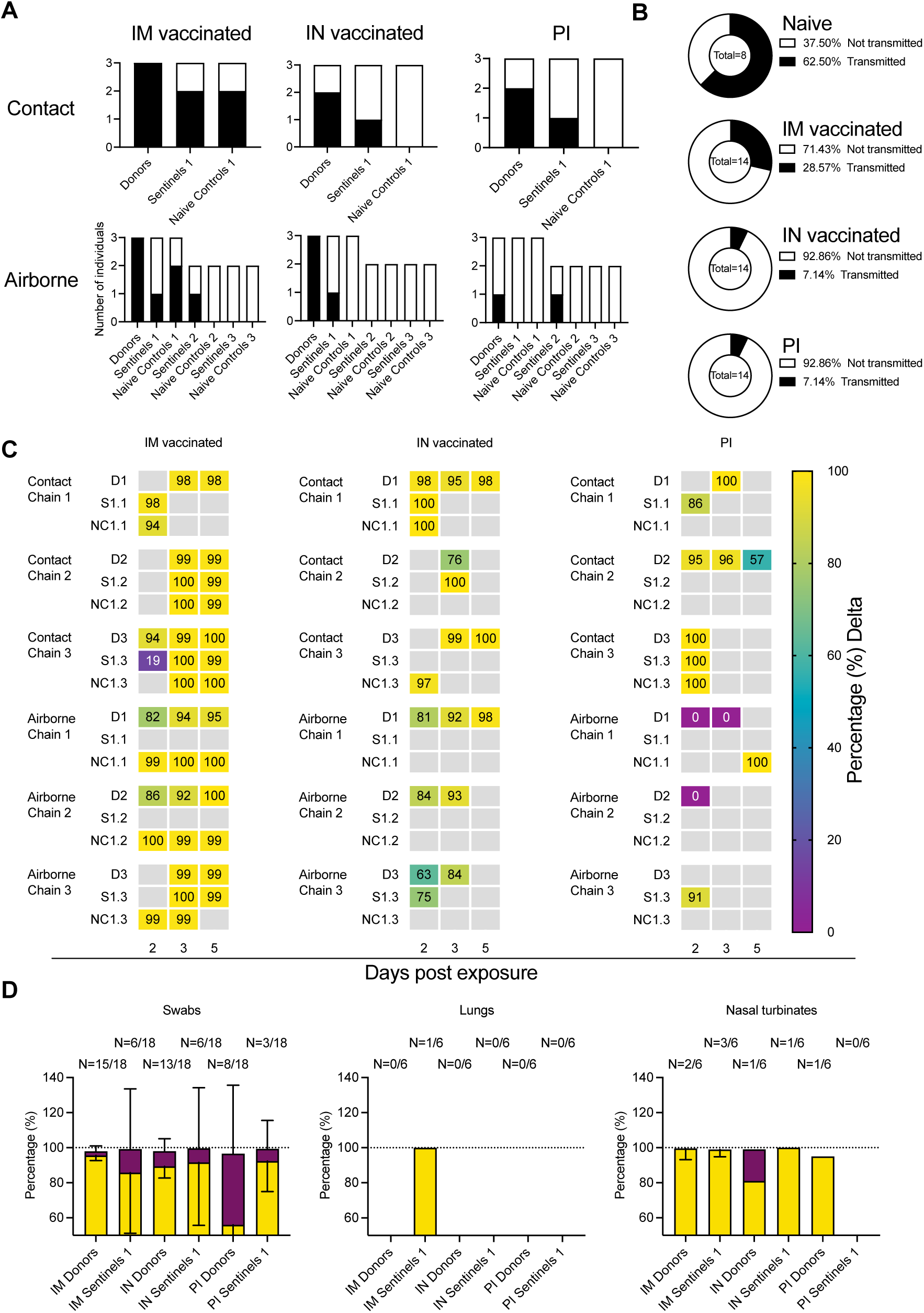
Transmission competitiveness of Delta and Omicron in animal groups with pre-existing immunity. Transmission efficiency and viral competitiveness in IN or IM vaccinated and PI hamsters. **A**. Summary of infection status for donors and sentinels. Oropharyngeal swabs were taken on 2, 3, and 5 DPI/DPE, and lungs and nasal turbinates collected at 5 DPI/DPE. Individuals were considered infected, if 2/5 samples were positive for sgRNA. Bar charts depict summary of individuals, divided by contact and airborne chains. **B**. Pie charts summarizing transmission efficiency between naïve, IM vaccinated, IN vaccinated, and PI hamsters across all airborne transmission events. Number of events is indicated within each pie chart. Colors refer to legends on right. **C**. The receptor binding domain of the SARS-CoV-2 spike was sequenced for all sgRNA positive swab samples taken on 2, 3, and 5 DPI/DPE. Heatmap displaying all sgRNA positive samples from each individual for each chain and showing percentage of Delta detected. Colors refer to legend on right (D = donor, S = sentinel, NC = naïve control), grey = no sgRNA present in the sample or sequencing unsuccessful. **D**. Overall percentage of Delta and Omicron in all sgRNA positive samples in each group, separated by sample type. Bar charts depicting mean and 95% CI. Number of sgRNA positive samples over all samples analyzed is indicated on top. Yellow = Delta, purple = Omicron.

Next, we compared the magnitude of overall shedding (AUC of sgRNA recovered in oral swabs on 2, 3, and 5 DPE) between naïve sentinels 1 exposed to naïve donors, and the IM, IN and PI sentinels 1 and their respective naïve controls (**Supplemental Figure 2 A**). IM vaccination showed the least effect on cumulative shedding compared to naïve sentinels, while IN vaccination and previous infection impacted cumulative shedding more. For IM vaccinated, IN vaccinated, and PI sentinels 1, cumulative shedding was significantly reduced compared to naïve sentinels 1 (p = 0.0374 (IM vaccinated), p = 0.0207 (IN vaccinated), and p = 0.0039 (PI), N = 6, two-way ANOVA, followed by Šídák’s multiple comparisons test). In contrast, while naïve controls shed similar amounts to naïve sentinels 1 in the IM group, we only observed significant reduction in cumulative shedding in naïve controls in the IN vaccinated group (p = 0.001) and the PI group (p = 0.004). When excluding all animals with no detectable sgRNA in any oral swab, the magnitude of cumulative shedding did not differ between sentinels with pre-existing immunity and their respective naïve controls. We observed a similar pattern when comparing lung pathology as measured by lung:body weight ratio (**Supplemental Figure 2 B**). Comparing sentinels 1, vaccination and previous infection offered significant protection (p = 0.0432 (IM vaccinated), p = 0.033 (IN vaccinated), and p = 0.002 (PI), N = 6, two-way ANOVA, followed by Šídák’s multiple comparisons test). Protection was also increased for naïve controls, but it was only significant in the PI group (p = 0.0238). We did not see a significant difference in the protection from lung pathology between sentinels with pre-existing immunity and their respective naïve controls.

### Existing immunity impacts Omicron intra- and inter-host competitiveness

To elucidate the intra- and inter-host competitiveness of Delta and Omicron, we determined the relative variant composition in sgRNA positive swabs and tissue samples by next-generation sequencing. In a few hamsters, Omicron was the dominant variant (**Supplementary Table 2**). Overall, Delta outcompeted Omicron in the directly infected donors and the sentinels across all groups (**Figure 5 C**). However, compared to the percentage of Omicron sequences in swab samples from the naïve animals (< 2%), Omicron was more prevalent in swab samples from hamsters with pre-existing immunity: Donors: IM vaccinated = 2.4%, IN vaccinated = 8.7%, and PI = 40.6%; Sentinels 1: IM vaccinated = 13.4 %, IN vaccinated = 8.0%, and PI = 6.9% (**Figure 5 D)**. This trend did not appear in tissue samples, and no Omicron was recovered in the nasal turbinates of either IM vaccinated or PI animals, with the exception of one IN donor (18% Omicron), nor in the lungs of IM vaccinated animals. No sgRNA was recovered from lungs of IN vaccinated or PI animals. These data suggest that immune pressure may be different between physiological compartments within the host, or that in the hamster model the initial relative advantage provided by pre-existing immunity is rapidly lost once infection is established.

### Intratracheal inoculation with Omicron leads to lung replication and pathology, but not increased transmission

Considering the difference in phenotype of Omicron compared to Delta in the Syrian hamster [22], we set out to better understand the mechanisms of transmission and pathogenesis for Omicron in hamsters. Omicron shows limited lower respiratory tract dissemination through the loss of TMPRSSII affinity due to mutations in the cleavage site [9, 23]. This could suggest that the route of administration is key to changing the transmission profile and inducing pathogenesis in the lower respiratory tract. We compared intratracheal (IT) inoculation with Omicron with the IN route. The shedding profile for both inoculation routes was identical (**Figure 6 A**). We found that after IT inoculation, more virus replication was observed in the lungs (median sgRNA copies/gram (Log_10_): IN = 0, IT = 9.358, p = 0.0022, Mann-Whitney test, N = 6), but not in the nasal turbinates (**Figure 6 B**). This was accompanied by a significant increase in lung pathology (**Figure 6 C**), measured by increased lung:body weight ratio at 5 DPI in the IT group (lung:body (%) = 0.8662 (IN) / 1.669 (IT), p = 0.0022, Mann-Whitney test, N = 6) and observable lung lesions by gross pathology (**Figure 6 D**). Histopathological lesions and SARS-CoV-2 NP immunoreactivity (p = 0.0022, Mann-Whitney test, N = 6) in the alveoli were consistent with what has previously been described for other SARS-CoV-2 variants [18, 19]. (**Figure 6 E-F**). We then assessed if the change in tropism by inoculation route translated into differential virus transmission dynamics. We exposed sentinels (1:1 ratio donor:sentinel) either by contact or by airborne exposure to IN or IT infected donors (N = 3 for each variation), for 48 h starting on 1 DPI. In the IN group, sgRNA was detectable in two contact sentinels and one air sentinel on both sampling days, and in one IN contact sentinel in only one sample (**Table 1)**. In the IT group, sgRNA was detectable in one contact sentinel on both days and one contact sentinel in only one sample (**Table 1)**. No positive samples were found for the air sentinels. All sentinels positive in swabs also seroconverted (**Table 1**). Interestingly, gRNA could only be recovered in air samples on day 1 and 2 of exposure if the donor animal was IN inoculated, but not IT inoculated (**Figure 6 G**). These data imply that upper respiratory tract replication may be required for transmissibility through air in this model (1 out of 3 sentinels positive for the IN group, as compared to 0 out of 3 for the IT group), but less so for contact (3 out of 3 sentinels positive for the IN group, as compared to 2 out of 3 for the IT group).

**Figure 6.**
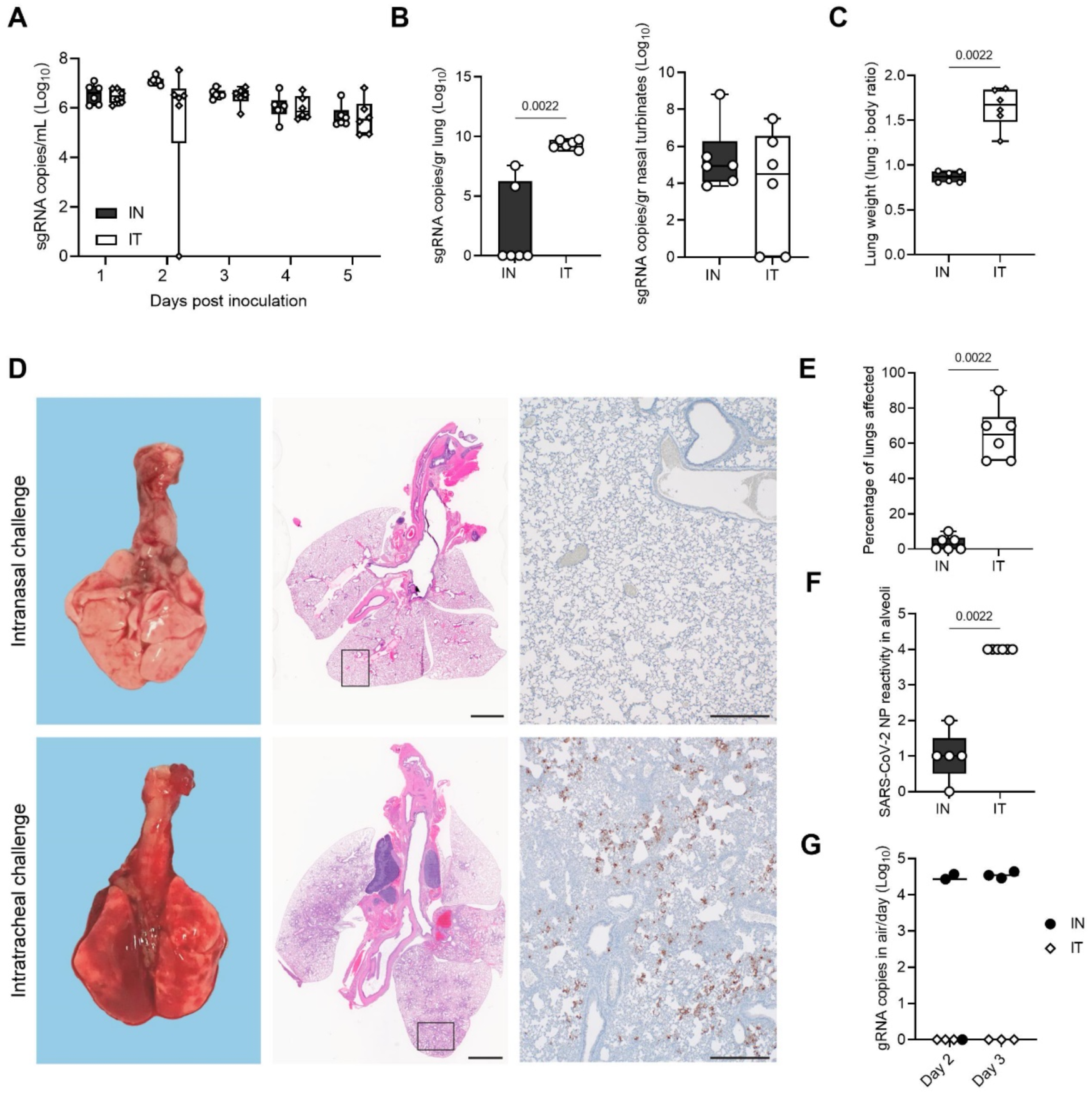
Recovery of lower respiratory tract replication and pathogenicity using intratracheal inoculation with Omicron. Syrian hamsters were inoculated with Omicron through the intranasal (IN) or intratracheal (IT) route (group size N = 6). Shedding and virus titers in tissues at 5 DPE were compared. **A**. Viral load as measured by sgRNA in oropharyngeal swabs collected at 1-5 days post inoculation. Whisker-plots depicting median, min and max values, and individual values, N = 6. **B**. sgRNA in lungs and nasal turbinates. Whisker-plots depicting median, min and max values, and individual values. Kruskal-Wallis test, N = 6. **C**. Lung weights (lung : body ratio). Whisker-plots depicting median, min and max values, and individual values, Kruskal-Wallis test, N = 6. **D**. Gross pathology of lungs on IN (top) and IT (bottom) inoculated animals at 5 DPE (left), histopathology (HE, middle), and immunohistochemistry against SARS-COV-2 NP (IHC, 200x, right) **E**. Percentage of lungs affected. **F**. Quantitative analysis of the NP reactivity. Whisker-plots depicting median, min and max values, and individual values. Kruskal-Wallis test, N = 6. **G**. For each airborne transmission, cage air was sampled in 24h intervals. Measurement of each individual cage is shown for gRNA. black = IN, white = IT. P-values stated were significant (<0.05).

**Table 1:** sgRNA shedding on 3 and 5 DPE and 14-day seroconversion of sentinel animals exposed to IN or IT inoculated donors. Shedding data is for N = 3 for contact and airborne transmission at 16.5 cm. Seroconversion of sentinels was measured by anti-spike SARS CoV-2 IgG ELISA and values are the average of two replicates, diluted 1:100. Cut-off = OD of 0.07 for positivity.

|  |  | Sentinel Shedding sgRNA<br>copies/mL (Log<sub>10</sub>) |  | Sentinel<br>seroconversion<br>(ELISA) |
| --- | --- | --- | --- | --- |
|  |  | Day 3 | Day 5 | Day 14 |
| IN Donor Contact | 1 | 0.00 | 6.71 | positive |
| IN Donor Contact | 2 | 5.14 | 5.88 | positive |
| IN Donor Contact | 3 | 5.06 | 5.72 | positive |
| IT Donor Contact | 7 | 6.26 | 5.16 | positive |
| IT Donor Contact | 8 | 0.00 | 0.00 | negative |
| IT Donor Contact | 9 | 6.74 | 0.00 | positive |
| IN Donor Air | 4 | 0.00 | 0.00 | negative |
| IN Donor Air | 5 | 0.00 | 0.00 | negative |
| IN Donor Air | 6 | 5.57 | 6.08 | positive |
| IT Donor Air | 10 | 0.00 | 0.00 | negative |
| IT Donor Air | 11 | 0.00 | 0.00 | negative |
| IT Donor Air | 12 | 0.00 | 0.00 | negative |

## Discussion

The ongoing circulation of SARS-CoV-2 VOCs and vaccinations have created a highly heterogeneous immune landscape in the human population. Household transmission analyses have revealed that vaccinations against SARS-CoV-2 can be effective in reducing transmission not only for SARS-CoV-2 Lineage A, but also VOCs including Delta [24]. Fully vaccinated and booster-vaccinated individuals are generally less susceptible to infection compared to unvaccinated individuals [25]. Vaccination may reduce pathogen load directly affecting the disease transmission dynamics. In experimental studies in the Syrian hamster, low heterologous vaccination-induced antibody titers were linked to a reduction in lower respiratory tract pathology and virus replication [26]. However, vaccine induced-SARS-CoV-2 immunity is typically not sterilizing [27],and transmission and virus replication in the upper respiratory tract are still observed after homologous or heterologous challenge. In addition, while the risk of reinfections in humans has been linked to the magnitude of clinical and serological presentation of the first infection, vaccinations have been found to reduce the risk of reinfection [28]. However, none of the currently licensed vaccines are able to completely block transmission. In particular, with the Omicron VOC, vaccine breakthrough and reinfections have frequently been reported. These are likely driven by a combination of waning immunity and antigenic drift [29]. There is a clear need for the development of vaccines with the potential to reduce upper respiratory tract replication and transmission while maintaining their ability to prevent lower respiratory tract disease.

AZD1222 is a replication-incompetent simian adenovirus–vectored vaccine encoding the Lineage A Spike (S) protein of Wuhan-1. Compared to IM vaccination, mucosal vaccination with the ChAdOx1 COVID19 vaccine (AZD1222) has been shown to be more efficient in preventing upper respiratory tract viral replication and shedding, while retaining the potential to prevent disease in pre-clinical models, including the Syrian hamster, ferrets, and rhesus macaques [20, 33]. We found here, that while the AZD1222 vaccine was based on the Lineage A S protein, both IM and IN vaccination provided protection from lung pathology after challenge with a Delta/Omicron mixture in the Syrian hamster. Our data supports increased protection of the lower respiratory tract after IN vaccination or previous infection compared to IM vaccination. As we and others previously demonstrated [20, 33], in this study the mucosal vaccination also decreased the viral load in the upper respiratory tract as compared to IM vaccination. However, in the PI animals only, did we observe a significant reduction in cumulative shedding compared to naïve hamsters.

Mucosal COVID-19 vaccines have been experimentally shown to reduce upper respiratory shedding, but not transmission, when assessed in a single contact transmission chain setting [30-32]. In addition, hamster studies have shown that previous infection protects against disease, but not upper respiratory tract replication, after homologous and heterologous reinfection [27, 34-36]. Similar dynamics were observed in our experimental setup. Vaccination did not completely block the transmission in the first round but, a disruption of the airborne transmission chain was achieved in the second iteration of the transmission chain. Our data show that vaccination resulted in markedly changed transmission and disease dynamics, and this effect was greater for IN than IM vaccination. Vaccination and PI also reduced the magnitude of shedding and disease severity in the lungs in sentinel animals as compared to sentinels exposed to naïve donors. This suggests that pre-existing immunity of the donor not only significantly reduces the likelihood of the first transmission event but also may impact the onwards transmission on a population level, magnifying the effect. The ability to block transmission appears to be related to the strength of the immune response, in which we observe a change in strength from IM to IN to PI. IN vaccination and PI were associated with higher binding antibody levels in serum and neutralizing titers as compared to IM vaccination, and IN vaccination reduced the potential for airborne transmission more robustly, to the same level as previous infection.

In this study, we did not observe differences in the magnitude of shedding found in naïve control sentinels and sentinels with pre-existing immunity after successful transmission from a vaccinated or PI donor. This might indicate that the transmission blocking efficacy of vaccinations may be a result mostly of reduced shedding of the donor as opposed to protection of the sentinel from infection, rather occurring on both ends of the transmission chain.

In humans, household transmission studies have demonstrated that immunity through vaccination does not provide equal protection across variants. Among vaccinated individuals, the Omicron VOC is generally 2.7-3.7 times more infectious than the Delta VOC [25]. However, this difference was absent among unvaccinated individuals, suggesting that increased transmissibility of the Omicron VOC is likely due to immune evasion [12, 37, 38]. In addition, more cases of household transmission from primary cases were observed with Omicron compared to Delta [39]. Whereas the antigenic differences between the original ancestral SARS-CoV-2 and most VOCs are relatively minimal, the exception is Omicron [12]. On average, a drop in neutralizing titers of ∼ 40 has been observed in sera from vaccinated and previously infected individuals [26, 40-42].

Omicron displays a reduced pathogenic phenotype in animal models. Virus replication in the lower respiratory tract is reduced both in rodents [43, 44] and non-human primates [45, 46]. Omicron also displays a reduced transmission efficiency compared to Lineage A, Alpha, and Delta in hamsters. Intratracheal inoculation with Omicron displayed a lung pathology and lower respiratory tract replication phenotype similar to that seen after intranasal inoculation with the other SARS-CoV-2 VOCs. Interestingly, contact transmission efficiency with Omicron was not reduced after intratracheal inoculation, and airborne transmission efficiency was not increased as compared to IN. These findings, combined with the absence of viral RNA in air samples taken from IT hamsters, suggest that replication in the upper respiratory tract is required for airborne transmission, but not for contact transmission.

Although Omicron showed reduced transmission potential, infection- or vaccine mediated immunity increased the relative transmission potential of Omicron compared to Delta. Delta out-competed Omicron in naïve hamster within and between hosts, suggesting overall greater fitness of Delta in that context. However, in PI animals with preexisting immunity, the relative frequency of Omicron increased compared to Delta. We also observed that the gap in neutralizing capacity between Omicron and Delta decreased more in the IN vaccinated and PI groups after challenge/re-infection with the Delta/Omicron mixture as compared to IM vaccinated animals. This could further suggest that in these groups, a replication advantage was present for Omicron initially, which led to increased antibody affinity maturation towards this variant. Our findings align with observations from another study where the authors showed that the presence of neutralizing antibodies against Delta, but not Omicron, could prevent Delta from outcompeting Omicron in hamsters [47]. This suggests, that even in hamsters, where Delta is intrinsically more transmissible, immune pressure can provide a direct advantage for antigenically-different viruses.

Our data demonstrate that pre-existing immunity and route of exposure directly influence disease manifestation and onwards transmission efficacy and potential. These data highlight the need to better understand SARS-CoV-2 transmission dynamics amidst the complexity of pre-existing immunity and the emergence of VOCs.

## Materials and Methods

### Ethics Statement

All animal experiments were conducted in an AAALAC International-accredited facility and were approved by the Rocky Mountain Laboratories Institutional Care and Use Committee following the guidelines put forth in the Guide for the Care and Use of Laboratory Animals 8th edition, the Animal Welfare Act, United States Department of Agriculture and the United States Public Health Service Policy on the Humane Care and Use of Laboratory Animals. Protocol number 2021-034-E. Work with infectious SARS-CoV-2 virus strains under BSL3 conditions was approved by the Institutional Biosafety Committee (IBC). For the removal of specimens from high containment areas, virus inactivation of all samples was performed according to IBC-approved standard operating procedures.

### Cells and viruses

The SARS-CoV-2 isolates used in this study are summarized in **Supplemental Table 3**. Virus propagation was performed in VeroE6 cells in DMEM supplemented with 2% fetal bovine serum, 1 mM L-glutamine, 50 U/mL penicillin and 50 μg/mL streptomycin (DMEM2). VeroE6 cells were maintained in DMEM supplemented with 10% fetal bovine serum, 1 mM L-glutamine, 50 U/mL penicillin and 50 μg/ml streptomycin. At regular intervals mycoplasma testing was performed. No mycoplasma or contaminants were detected. All virus stocks were sequenced; and no SNPs compared to the patient sample sequence were detected.

### Pseudotype entry assay

The spike coding sequences for SARS-CoV-2 variant Lineage A, Delta, and Omicron (MN985325, EPI_ISL_2441471, EPI_ISL_6699767, respectively) were truncated by deleting 19 aa at the C-terminus. The spike (S) proteins with the 19 aa deletions of coronaviruses were previously reported to show increased efficiency regarding incorporation into virions of VSV [48, 49]. These sequences were codon optimized for human cells, then appended with a 5′ kozak expression sequence (GCCACC) and 3′ tetra-glycine linker followed by nucleotides encoding a FLAG-tag sequence (DYKDDDDK). These spike sequences were synthesized and cloned into pcDNA3.1^+^(GenScript). Human and hamster ACE2 (Q9BYF1.2 and GQ262794.1, respectively), were synthesized and cloned into pcDNA3.1^+^ (GenScript). All DNA constructs were verified by Sanger sequencing (ACGT). BHK cells were seeded in black 96-well plates and transfected the next day with 100 ng plasmid DNA encoding human or hamster ACE2, using polyethylenimine (Polysciences). All downstream experiments were performed 24 h post-transfection. Pseudotype production was carried as previously described [50]. Briefly, plates pre-coated with poly-L-lysine (Sigma–Aldrich) were seeded with 293T cells and transfected the following day with 1,200 ng of empty plasmid and 400 ng of plasmid encoding coronavirus spike or no-spike plasmid control (green fluorescent protein (GFP)). After 24 h, transfected cells were infected with VSVΔG seed particles pseudotyped with VSV-G, as previously described [50, 51]. After one hour of incubating with intermittent shaking at 37 °C, cells were washed four times and incubated in 2 mL DMEM supplemented with 2% FBS, penicillin/streptomycin and L-glutamine for 48 h. Supernatants were collected, centrifuged at 500 x *g* for 5 min, aliquoted, and stored at −80 °C. BHK cells previously transfected with ACE2 plasmid of interest were inoculated with equivalent volumes of pseudotype stocks. Plates were then centrifuged at 1200 x *g* at 4 °C for one hour and incubated overnight at 37 °C. Approximately 18– 20 h post-infection, Bright-Glo luciferase reagent (Promega) was added to each well, 1:1, and luciferase was measured. Relative entry was calculated normalizing the relative light unit for spike pseudotypes to the plate relative light unit average for the no-spike control. Each figure shows the data for two technical replicates.

### Structural interaction analysis

The locations of the described spike mutations in the Delta and Omicron VOCs were highlighted on the SARS-CoV-2 spike structure (PDB 6ZGE [52]). To visualize the molecular interactions at the RBD – ACE2 binding interface, the crystal structure of the Alpha variant RBD and human ACE2 complex (PDB 7EKF [53]) was utilized. All figures were generated using The PyMOL Molecular Graphics System (https://www.schrodinger.com/pymol).

### VOC virus shedding comparison

For the comparison of virus shedding of all VOCs, four-to-six-week-old Syrian golden hamsters (N = 6 per group, Envigo Indianapolis) were inoculated intranasally with 40 μL containing 1 × 10^3^ TCID_50_ virus in sterile DMEM. Oropharyngeal swabs were collected in 1 mL of DMEM2 on day post infection 1-7.

### Variant transmission competitiveness between pre-existing immunity groups

Four-to-six-week-old female and male Syrian hamsters (ENVIGO) were used. Hamsters were randomly assigned to one of four groups: Naïve group, intramuscularly (IM) vaccinated group, intranasally (IN) vaccinated group, and PI (PI) group. For the IM vaccinated group, 16 animals received vaccine AZD1222 (2.5 × 10^8^ IU/animal) intramuscularly to two sites using a 25-gauge needle with a maximum injection volume of 200 μL. For the IN vaccinated group, 16 animals received vaccine AZD1222 (2.5 × 10^8^ IU/animal intranasally with a maximum injection volume of 60 μL. For the PI group, 16 naïve animals were exposed to Delta infected animals in direct contact over multiple days. Four hamsters were inoculated via the intranasal route with a total maximum dose of 10^4^ TCID_50_ SARS-CoV-2 Delta VOC. One infected hamster was co-housed with four naïve animals to allow for contact transmission to occur (ratio 1:4). 21 days post vaccination or challenge blood was collected for serology.

The transmission chains were conducted at least 28 days post vaccination or previous infection. Naïve controls were age matched. Naïve group: Donor hamsters (N = 6) were infected intranasally as described above with 1 × 10^4^ TCID_50_ SARS-CoV-2 at a 1:1 ratio of Omicron and Delta and individually housed. After 24 hours, three donor animals were placed into a new rodent cage and three donors were placed into the donor cage of an airborne transmission set-up of 16.5 cm distance at an airflow of 30 cage changes/h as described by Port et al. [14]. Sentinels (sentinels 1, N = 3) were placed into either the same cage (contact, N = 3,1:1 ratio) or the sentinel cage of the airborne transmission caging (airborne, N = 3, 1:1 ratio). Hamsters were co-housed for 48 h. Donor animals were re-housed into regular rodent caging and sentinels 1 were placed into either a new rodent cage or the donor cage of a new airborne transmission set-up. New sentinels (sentinels 2, N = 3 for contact and N = 3 for airborne) were placed into the same new rodent cage or the sentinel cage of the airborne transmission caging (1:1) at 16.5 cm distance at an airflow of 30 changes/h. Hamsters were co-housed for 48 h. Sentinels 1 were then re-housed into regular rodent caging and N = 4/6 sentinels 2 were placed into either a new rodent cage or the donor cage of a new airborne transmission set-up. New sentinels (sentinels 2, N = 2 for contact and N = 2 for airborne) were placed into the same new rodent cage or the sentinel cage of the airborne transmission caging (1:1) at 16.5 cm distance at an airflow of 30 changes/h. Hamsters were co-housed for 72 h. Then all were re-housed to regular rodent caging and monitored until 5 DPE.

#### Vaccinated groups

Donor hamsters (N = 6 for IM and IN vaccination, respectively) were infected intranasally as described above with 1 × 10^4^ TCID_50_ SARS-CoV-2 at a 1:1 ratio of Omicron and Delta and individually housed. After 24 hours for each group, three donor animals were placed into a new rodent cage and three donors were placed into the donor cage of an airborne transmission set-up. Equally vaccinated sentinels (sentinels 1) and completely naïve animals (naïve Controls) were placed into either the same cage (contact, N = 3,1:2 ratio) or the sentinel cage of the airborne transmission caging (airborne, N = 3, 1:2 ratio). Hamsters were co-housed for 48 h. Donor animals and sentinels were re-housed into regular rodent caging and monitored until 5 DPE.

#### PI group

Donor hamsters (N = 6) were infected intranasally as described above with 1 × 10^4^ TCID_50_ SARS-CoV-2 at a 1:1 ratio of Omicron and Delta and individually housed. After 24 hours, three donor animals were placed into a new rodent cage and three donors were placed into the donor cage of an airborne transmission set-up. Equally PI sentinels (sentinels 1) and completely naïve animals (naïve controls) were placed into either the same cage (contact, N = 3,1:2 ratio) or the sentinel cage of the airborne transmission caging (airborne, N = 3, 1:2 ratio). Hamsters were co-housed for 48 h. Donor animals and sentinels were re-housed into regular rodent caging and monitored until 5 DPE. Oropharyngeal swabs were taken for all animals at 2, 3, and 5 DPI/DPE. All animals were euthanized at 5 DPI/DPE for collection of lung tissue and nasal turbinates and serum. To ensure no cross-contamination, the donor cages and the sentinel cages were never opened at the same time, sentinel hamsters were not exposed to the same handling equipment as donors, and the equipment was disinfected with either 70% ETOH or 5% Microchem after each sentinel. Regular bedding was replaced by alpha-dri bedding to avoid the generation of dust particles.

### Comparison between intranasal and intratracheal inoculation

Four-to-six-week-old male Syrian hamsters (ENVIGO) were used. Animals were randomly assigned to two groups, intratracheal and intranasal inoculation, and inoculated with 1 × 10^4^ TCID_50_ SARS-CoV-2 in a volume of 40 μL (IN) or 100μL (IT) (N = 6). Animals were then individually housed, swabbed daily in the oropharyngeal cavity, and lungs and nasal turbinates collected at day 5. On day 1, each animal was either co-housed with a naïve sentinel (contact, N = 3) or placed into the upstream cage of a short-distance aerosol transmission cage (16.5cm) and one sentinel placed adjacent (air, N = 3). Animals were exposed at a 1:1 ratio, for 48 hours. Air was sampled in 24h intervals for the air transmission set-ups as described previously [14]. Sentinels were swabbed on days 3 and 5 post exposure start, and serum collected on day 14.

### Viral RNA detection

Swabs from hamsters were collected as described above. Then, 140 μL was utilized for RNA extraction using the QIAamp Viral RNA Kit (Qiagen) using QIAcube HT automated system (Qiagen) according to the manufacturer’s instructions with an elution volume of 150 μL. For tissues, RNA was isolated using the RNeasy Mini kit (Qiagen) according to the manufacturer’s instructions and eluted in 60 μL. Sub-genomic (sg) and genomic (g) viral RNA was detected by qRT-PCR [54, 55]. RNA was tested with TaqMan™ Fast Virus One-Step Master Mix (Applied Biosystems) using QuantStudio 3 Flex Real-Time PCR System (Applied Biosystems). SARS-CoV-2 standards with known copy numbers were used to construct a standard curve and calculate copy numbers/mL or copy numbers/g. The detection limit for the assay was 10 copies/reaction, and samples below this limit were considered negative.

### Virus titration

Viable virus in tissue samples was determined as previously described [56]. In brief, lung tissue samples were weighed, then homogenized, in 1 mL of DMEM (2% FBS). Swabs were used undiluted. VeroE6 cells were inoculated with ten-fold serial dilutions of homogenate, incubated 1 hours at 37°C, and the first two dilutions washed twice with 2% DMEM. For swab samples, cells were inoculated with ten-fold serial dilutions and no wash was performed. After 6 days, cells were scored for cytopathic effect. TCID_50_/mL was calculated by the method of Spearman-Karber. To determine titers in air samples, a plaque assay was used. VeroE6 cells were inoculated with 200 μL/well (48-well plate) of undiluted samples, with no wash performed. Plates were spun for 1 hour at room temperature at 1000 rpm. 800 uL of CMC (500 mL MEM (Cat#10370, Gibco, must contain NEAA), 5 mL PenStrep, 7.5 g carboxymethylcellulose (CMC, Cat# C4888, Sigma, sterilize in autoclave) overlay medium was added to each well and plates incubated for 6-days at 37°C. Plates were fixed with 10% formalin overnight, then rinsed and stained with 1% crystal violet for 10 min. Plaques were counted.

### ELISA

Serum samples were analyzed as previously described [57]. In brief, maxisorp plates (Nunc) were coated with 50 ng spike protein (generated in-house) per well. Plates were incubated overnight at 4°C. Plates were blocked with casein in phosphate buffered saline (PBS) (ThermoFisher) for 1 hour at room temperature. Serum was diluted 2-fold in blocking buffer and samples (duplicate) were incubated for 1 hour at room temperature. Secondary goat anti-hamster IgG Fc (horseradish peroxidase (HRP)-conjugated, Abcam) spike-specific antibodies were used for detection and visualized with KPL TMB 2-component peroxidase substrate kit (SeraCare, 5120-0047). The reaction was stopped with KPL stop solution (Seracare) and plates were read at 450 nm. The threshold for positivity was calculated as the average plus 3 x the standard deviation of negative control hamster sera.

### MESO QuickPlex Assay

The V-PLEX SARS-CoV-2 Panel 23 (IgG) kit from Meso Scale Discovery was used to test binding antibodies against the spike protein of the different SARS-CoV-2 VOCs, with serum obtained from hamsters 14 DPI diluted at 10,000X. A standard curve of pooled hamster sera positive for SARS-CoV-2 spike protein was serially diluted 4-fold. To prepare a secondary antibody, a goat anti-hamster IgG cross-adsorbed secondary antibody (ThermoFisher) was conjugated using the MSD GOLD SULFO-TAG NHS-Ester Conjugation Pack (MSD). The secondary antibody was diluted 10,000X. The plates were prepped, and samples were run according to the kit’s instruction manual. After the plates were read by the MSD instrument, data was analyzed with the MSD Discovery Workbench Application.

### Virus neutralization

Heat-inactivated γ-irradiated sera were two-fold serially diluted in DMEM. 100 TCID_50_ of SARS-CoV-2 were added. After 1 hour of incubation at 37°C and 5% CO_2_, the virus:serum mixture was added to VeroE6 cells. CPE was scored after 5 days at 37 °C and 5% CO_2_. The virus neutralization titer was expressed as the reciprocal value of the highest dilution of the serum which still inhibited virus replication.

### Next-generation sequencing of virus

Total RNA was extracted from oral swabs, lungs, and nasal turbinates using the Qia Amp Viral kit (Qiagen, Germantown, MD), eluted in EB, and viral Ct values were calculated using real-timePCR. Subsequently, 11 μL of extracted RNA were used as template in the ARTIC nCoV-2019 sequencing protocol V.1 (Protocols.io - https://www.protocols.io/view/ncov-2019-sequencing-protocol-bbmuik6w) to generate first-strand cDNA. Five microliters were used as template for Q5 HotStart Polymerase PCR (Thermo Fisher Sci, Waltham, MA) together with 10 uM stock of a single primer pair from the ARTIC nCoV-2019 v3 Panel (Integrated DNA Technologies, Belgium); specifically, 76L_alt3 and 76R_alt0. Following 35 cycles and 55°C annealing temperature, products were AmPure XP cleaned and quantitated with Qubit (Thermo Fisher Sci) fluorometric quantitation as per instructions. Following visual assessment of 1 μL on a Tape Station D1000 (Agilent Technologies, Santa Clara, CA), a total of 400 ng of product was taken directly into TruSeq DNA PCR-Free Library Preparation Guide, Revision D. (Illumina, San Diego, CA) beginning with the Repair Ends step (q.s. to 60 μL with RSB). Subsequent clean-up consisted of a single 1:1 AmPure XP/reaction ratio, and all steps followed the manufacturer’s instructions including the Illumina TruSeq CD (96) indexes. Final libraries were visualized on a BioAnalyzer HS chip (Agilent Technologies) and quantified using KAPA Library Quant Kit - Illumina Universal qPCR Mix (Kapa Biosystems, Wilmington, MA) on a CFX96 Real-Time System (BioRad, Hercules, CA). Libraries were diluted to 2 nM stock, pooled together in equimolar concentrations, and sequenced on the Illumina MiSeq instrument (Illumina) as paired-end 2 × 250 base pair reads. Because of the limited diversity of a single-amplicon library, 20% PhiX was added to the final sequencing pool to aid in final sequence quality. Raw fastq reads were trimmed of Illumina adapter sequences using cutadapt version 1.1227, and then trimmed and filtered for quality using the FASTX-Toolkit (Hannon Lab, CSHL). To process the ARTIC data, a custom pipeline was developed [58]. Fastq read pairs were first compared to a database of ARTIC primer pairs to identify read pairs that had correct, matching primers on each end. Once identified, the ARTIC primer sequence was trimmed off. Read pairs that did not have the correct ARTIC primer pairs were discarded. Remaining read pairs were collapsed into one sequence using AdapterRemoval [59] requiring a minimum 25 base overlap and 300 base minimum length, generating ARTIC amplicon sequences. Identical amplicon sequences were removed, and the unique amplicon sequences were then mapped to the SARS-CoV-2 genome (MN985325.1) using Bowtie2 [60]. Aligned SAM files were converted to BAM format, then sorted and indexed using SAMtools [61]. Variant calling was performed using Genome Analysis Toolkit (GATK, version 4.1.2) HaplotypeCaller with ploidy set to 2 [62]. Single nucleotide polymorphic variants were filtered for QUAL > 200 and quality by depth (QD) > 20 and indels were filtered for QUAL > 500 and QD > 20 using the filter tool in bcftools, v1.9 [61].

### Histopathology

Necropsies and tissue sampling were performed according to IBC-approved protocols. Tissues were fixed for a minimum of 7 days in 10% neutral buffered formalin with 2 changes. Tissues were placed in cassettes and processed with a Sakura VIP-6 Tissue Tek, on a 12-hour automated schedule, using a graded series of ethanol, xylene, and PureAffin. Prior to staining, embedded tissues were sectioned at 5 μm and dried overnight at 42°C. Using GenScript U864YFA140-4/CB2093 NP-1 (1:1000) specific anti-CoV immunoreactivity was detected using the Vector Laboratories ImPress VR anti-rabbit IgG polymer (# MP-6401) as secondary antibody. The tissues were then processed using the Discovery Ultra automated processor (Ventana Medical Systems) with a ChromoMap DAB kit Roche Tissue Diagnostics (#760-159). Anti-CD3 immunoreactivity was detected utilizing a primary antibody from Roche Tissue Diagnostics predilute (#790-4341), secondary antibody from Vector Laboratories ImPress VR anti-rabbit IgG polymer (# MP-6401) and visualized using the ChromoMap DAB kit from Roche Tissue Diagnostics (#760-159). Anti-PAX5 immunoreactivity was detected utilizing a primary antibody from Novus Biologicals at 1:500 (#NBP2-38790), secondary antibody from Vector Laboratories ImPress VR anti-rabbit IgG polymer (# MP-6401) and visualized using the ChromoMap DAB kit from Roche Tissue Diagnostics (#760-159).

### Morphometric analysis

CD3 and PAX5 IHC stained sections were scanned with an Aperio ScanScope XT (Aperio Technologies, Inc., Vista, CA) and analyzed using the ImageScope Positive Pixel Count algorithm (version 9.1). The default parameters of the Positive Pixel Count (hue of 0.1 and width of 0.5) detected antigen adequately.

## Acknowledgements

We would like to thank Bob Fischer and Shane Gallogly for help with experiments. We thank Tina Thomas, Rebecca Rosenke, and Dan Long for assistance with histology; RMVB animal care staff for taking care of the animals. The following reagent was obtained through: CDC: SARS-CoV-2/human/USA/WA-CDC-WA1/2020, Lineage A. BEI Resources, NIAID, NIH: SARS-CoV-2 variant Alpha (B.1.1.7) (hCoV320 19/England/204820464/2020, EPI_ISL_683466) and variant Delta (B.1.617.2/) (hCoV-19/USA/KY-CDC-2-4242084/2021, EPI_ISL_1823618). Variant Beta (B.1.351) isolate name: hCoV-19/USA/MD-HP01542/2021, EPI_ISL_890360, and variant Gamma (P.1) isolate name: hCoV-19/USA/MD-HP03867/2021, EPI_ISL_1468644, were contributed by Johns Hopkins Bloomberg School of Public Health: Andrew Pekosz. Variant Omicron (B.1.1.529. BA.1) isolate name: hCoV-19/USA/GA-EHC-2811C/2021, EPI_ISL_7171744, was contributed by Mehul Suthar. We thank Andrew Pekosz and Mehul Suthar for gracefully sharing viruses.

## Funding

This work was supported by the Intramural Research Program of the National Institute of Allergy and Infectious Diseases (NIAID), National Institutes of Health (NIH) (1ZIAAI001179-01). This work was part of NIAID’s SARS-CoV-2 Assessment of Viral Evolution (SAVE) Program.

## Notes

### Competing Interest Statement

The authors have declared no competing interest.

