## Supplemental Files for "Infection- or vaccine mediated immunity reduces SARS-CoV-2 transmission, but increases competitiveness of Omicron in hamsters"

* These first authors contributed equally

### These senior authors contributed equally

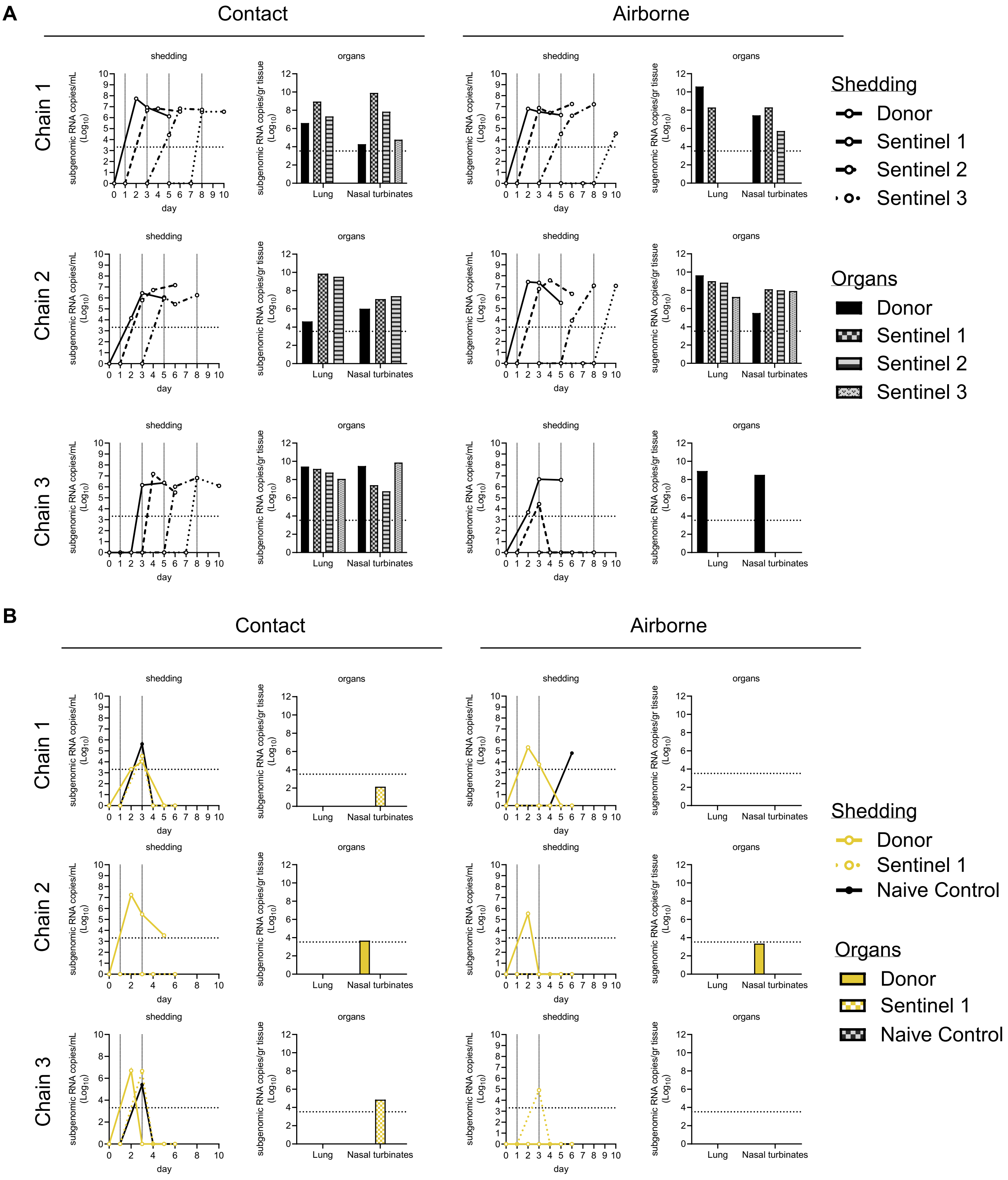
A
Contact
Airborne
Chain 1
Chain 2
Chain 3
shedding
organs
subgenomic RNA copies/mL (Log10)
subgenomic RNA copies/gr tissue (Log10)
sugenomic RNA copies/gr tissue (Log10)
day
Lung
Nasal turbinates
Shedding
Donor
Sentinel 1
Sentinel 2
Sentinel 3
Organs
Donor
Sentinel 1
Sentinel 2
Sentinel 3
B
Contact
Airborne
Chain 1
Chain 2
Chain 3
Shedding
Donor
Sentinel 1
Naive Control
Organs
Donor
Sentinel 1
Naive Control

C

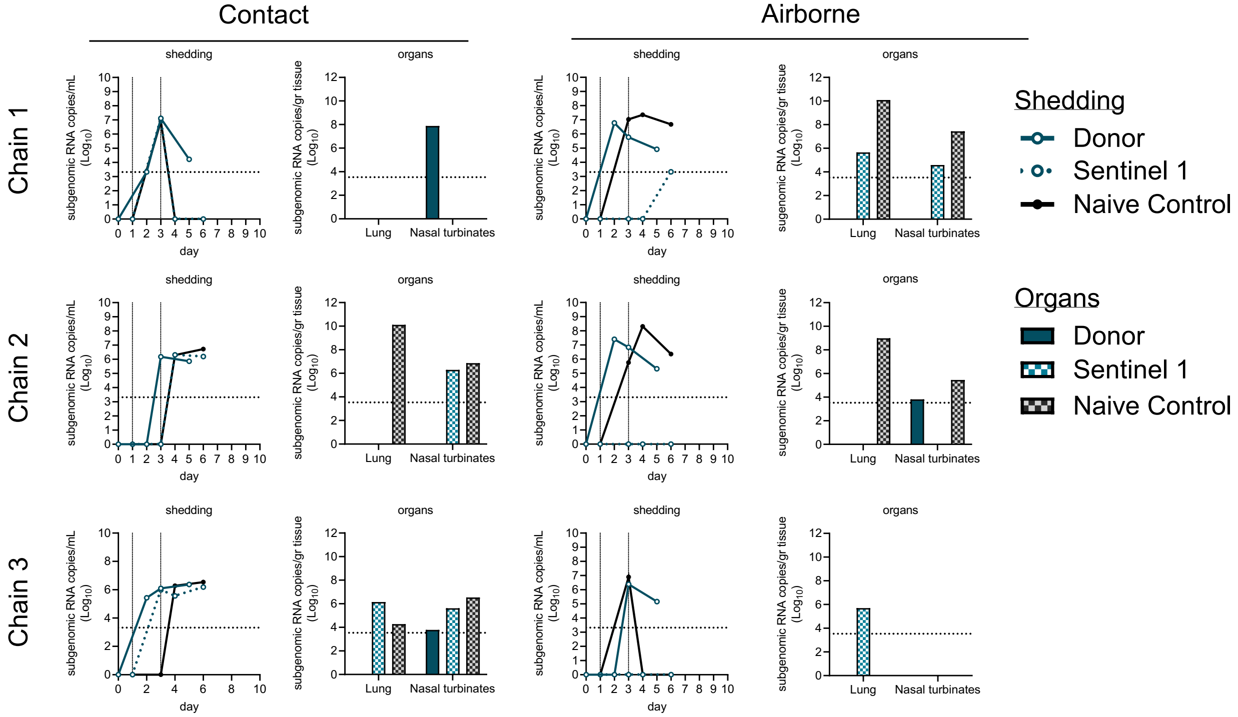
Contact
Airborne
Chain 1
Chain 2
Chain 3
shedding
organs
subgenomic RNA copies/mL (Log10)
subgenomic RNA copies/gr tissue (Log10)
sugenomic RNA copies/gr tissue (Log10)
day
Lung
Nasal turbinates
Shedding
Donor
Sentinel 1
Naive Control
Organs
Donor
Sentinel 1
Naive Control

D

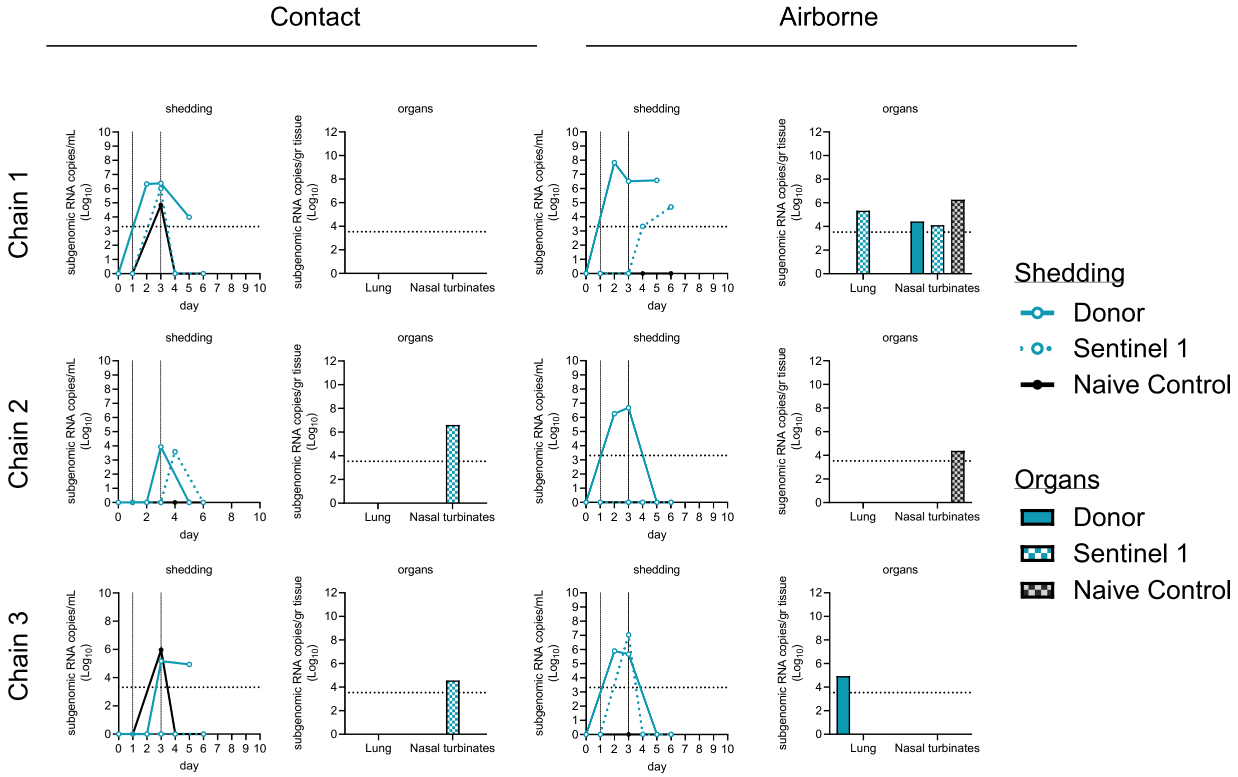
Contact
Airborne
Chain 1
Chain 2
Chain 3
shedding
organs
subgenomic RNA copies/mL (Log10)
subgenomic RNA copies/gr tissue (Log10)
sugenomic RNA copies/gr tissue (Log10)
day
Lung
Nasal turbinates
Shedding
Donor
Sentinel 1
Naive Control
Organs
Donor
Sentinel 1
Naive Control

**Supplemental Figure 1. Shedding and organ titers of individual animals in each chain.** Animals were grouped into either airborne or contact transmission chains by immune state (N = 3). Chains consisted of either a donor and three consecutive sentinels (naïve chains) or a donor and two sentinels, one with previous immunity and one naïve control (IM, IN, and PI chains). Donors were infected with 1 x $10^4$ $TCID_{50}$ SARS-CoV-2 at a 1:1 ratio of Omicron and Delta and 24 hours later exposed to either a sentinel and naïve animal (IM, IN, and PI chains) or to the first sentinel (naïve chains) for 48 hours. Naïve chains continued in a similar fashion over four total animals. Replicating virus was measured via oropharyngeal swabs at 2, 3, and 5 DPI/DPE, and in lung and nasal turbinate samples at 5 DPI/DPE. Viral shedding is depicted in oropharyngeal swabs and viral replication in lung and nasal turbinate samples of all animals in contact (left) and airborne (right) chains. Viral shedding and replication are displayed as $log_{10}$ (sgRNA copies/mL) in swabs and $log_{10}$ (sgRNA copies/gram of tissue) in tissues. Horizontal lines indicate limit of detection. Vertical lines indicate exposure events in the chain. **A.** Naïve chains. **B.** Previously infected chains, **C.** Intramuscularly vaccinated chains. **D.** Intranasally vaccinated chains. Colours refer to legends on the right.

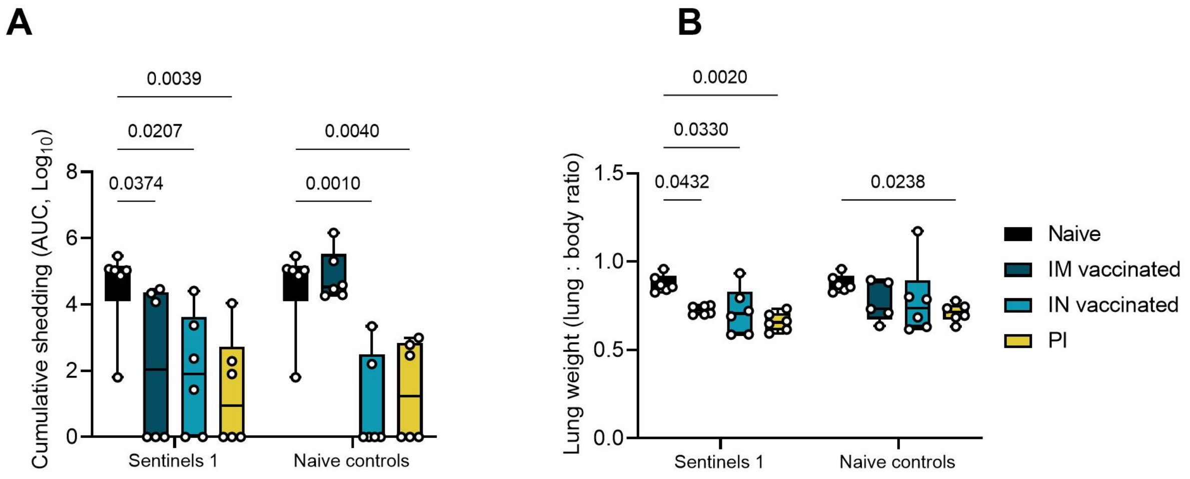


**Supplemental Figure 2. Reduction of disease severity and shedding in sentinels. A.** Cumulative shedding. Area under the curve (AUC) of sgRNA measured in oral swabs taken on 2, 3, and 5 DPE. Whisker-plots depicting median, min and max values, and individual values, N = 6, ordinary two-way ANOVA, followed by Šídák's multiple comparisons test. **B.** Lung samples were collected at 5 DPE for sentinels and naïve controls. Lung weights (lung:body weight ratio). Whisker-plots depicting median, min and max values, and individual values, N = 6, ordinary two-way ANOVA, followed by Šídák's multiple comparisons test. Black = naïve, dark blue = IM vaccinated, light blue = IN vaccinated, yellow = PI. P-values stated were significant (<0.05). Naïve sentinels 1 are depicted twice in each graph for visualization purposes only.

**Supplemental Table 1:** Pathological Assessment of IN, IM vaccinated or PI Syrian hamsters on day 5 post challenge. nsf = no significant findings. y = yes. n = no.

| Group | Naïve | | | | | | IM vaccinated | | | | | | IN vaccinated | | | | | | Previously infected | | | | | |
|---|---|---|---|---|---|---|---|---|---|---|---|---|---|---|---|---|---|---|---|---|---|---|---|---|
| Lesions visible grossly | y | y | n | y | y | y | y | y | y | ? | y | n | n | n | n | N | n | n | n | n | ? | n | n | n |
| Lung | | | nsf | | | | | | | nsf | | | nsf | nsf | nsf | nsf | | nsf | | nsf | nsf | nsf | nsf | |
| % total area affected | 60 | 60 | | 60 | 70 | 80 | 30 | <10 | 20 | | 20 | <10 | | | | | 10 | | <10 | | | | | <10 |
| Interstitial Pneumonia | | | | | | | | | | | | | | | | | | | | | | | | |
| Syncytial Cell | y | y | n | y | y | y | y | n | n | n | n | n | n | n | n | n | n | n | n | n | n | n | n | n |
| Alveolar inflammation or exudate | y | y | n | y | y | y | y | y | y | n | y | y | n | n | n | n | n | n | n | n | n | n | n | n |
| Endotheleitis, vasculitis | y | y | n | y | y | y | y | y | y | n | y | n | n | n | n | n | n | n | n | n | n | n | n | n |
| Type II pneumocyte hyperplasia | n | n | n | n | n | n | y | n | n | n | n | n | n | n | n | n | n | n | n | n | n | n | n | n |
| Hemorrhage, fibrin and or edema | y | y | n | y | y | y | n | n | y | n | n | n | n | n | n | n | n | n | n | n | n | n | n | n |
| Bronchiolar hyperplasia | y | y | n | y | y | y | y | n | n | n | n | n | n | n | n | n | n | n | n | n | n | n | n | n |
| SARS-CoV-2 IHC bronchioles | 3 | 2 | 0 | 2 | 2 | 2 | 0 | 0 | 3 | 0 | 0 | 1 | 0 | 0 | 0 | 0 | 0 | 0 | 0 | 0 | 0 | 0 | 0 | 0 |
| SARS-CoV-2 IHC alveoli | 4 | 3 | 0 | 3 | 4 | 4 | 1 | 0 | 3 | 0 | 0 | 0 | 0 | 0 | 0 | 0 | 0 | 0 | 0 | 0 | 0 | 0 | 0 | 0 |
| Positive pixel analysis CD3 | 16 | 17 | 7 | 14 | 14 | 16 | 6 | 10 | 10 | 9 | 9 | 10 | 7 | 8 | 8 | 8 | 6 | 9 | 8 | 7 | 8 | 7 | 8 | 6 |
| Positive pixel analysis PAX5 | 3 | 2 | 2 | 2 | 2 | 2 | 1 | 2 | 2 | 2 | 2 | 2 | 2 | 2 | 2 | 2 | 2 | 2 | 1 | 1 | 2 | 2 | 2 | 2 |

**Supplemental Table 2:** Shedding and tissue titers for each transmission chain. IM = intramuscularly vaccinated, IN = intranasally vaccinated, PI = Previously infected, BDL = Below qRT-PCR detection limit, sgRNA = sub-genomic RNA. Swab days = 1, 3 and 5 days

| | | | | sgRNA Ct-value | | | | | % Delta reads | | | | | % Omicron reads | | | | |
|---|---|---|---|---|---|---|---|---|---|---|---|---|---|---|---|---|---|---|
| **Transmission route** | **Chain** | **Chain link** | **Immune status** | **Day 2 swab** | **Day 3 swab** | **Day 5 swab** | **Lung** | **Nasal turbinates** | **Day 2 swab** | **Day 3 swab** | **Day 5 swab** | **Lung** | **Nasal turbinates** | **Day 2 swab** | **Day 3 swab** | **Day 5 swab** | **Lung** | **Nasal turbinates** |
| air | 1 | Donor | IM | 25 | 28 | 32 | BDL | BDL | 82 | 94 | 95 | | | 17 | 0 | 0 | | |
| air | 1 | Naïve Control | IM | 24 | 23 | 26 | 15 | 23 | 99 | 100 | 100 | 100 | 100 | 0 | 0 | 0 | 0 | 0 |
| air | 1 | Sentinel 1 | IM | BDL | BDL | 37 | 30 | 32 | | | | | 97 | | | | | 0 |
| contact | 1 | Donor | IM | 40 | 24 | 34 | BDL | 22 | | 98 | 98 | | 99 | 0 | 0 | 0 | | 0 |
| contact | 1 | Naïve Control | IM | 25 | BDL | BDL | BDL | BDL | 94 | | | | | 0 | | | | |
| contact | 1 | Sentinel 1 | IM | 24 | BDL | BDL | BDL | BDL | 98 | | | | | 0 | | | | |
| air | 2 | Donor | IM | 23 | 25 | 30 | BDL | 35 | 86 | 92 | 100 | | 100 | 13 | 7 | 0 | | 0 |
| air | 2 | Naïve Control | IM | 28 | 20 | 27 | 19 | 30 | 100 | 99 | 99 | 100 | 100 | 0 | 0 | 0 | 0 | 0 |
| air | 2 | Sentinel 1 | IM | BDL | BDL | BDL | BDL | BDL | | | | | | | | | | |
| contact | 2 | Donor | IM | BDL | 27 | 28 | BDL | BDL | | 99 | 99 | | | | 0 | 0 | 0 | |
| contact | 2 | Naïve Control | IM | BDL | 27 | 25 | 16 | 26 | | 100 | 99 | 100 | 100 | | 0 | 0 | 0 | 0 |
| contact | 2 | Sentinel 1 | IM | BDL | 27 | 27 | BDL | 28 | | 100 | 99 | | 100 | | 0 | 0 | | 0 |
| air | 3 | Donor | IM | BDL | 26 | 31 | BDL | BDL | | 99 | 99 | | | | 0 | 0 | | |
| air | 3 | Naïve Control | IM | 25 | BDL | BDL | BDL | BDL | 99 | | | | | 0 | 0 | | | |
| air | 3 | Sentinel 1 | IM | BDL | BDL | BDL | 28 | BDL | | | | | | | | | | |
| contact | 3 | Donor | IM | 30 | 27 | 27 | BDL | 35 | 94 | 99 | 100 | | | 0 | 0 | 0 | | 0 |
| contact | 3 | Naïve Control | IM | BDL | 27 | 26 | 34 | 31 | | 100 | 100 | | 100 | | 0 | 0 | | 0 |
| contact | 3 | Sentinel 1 | IM | 28 | 29 | 27 | 27 | 29 | 19 | 100 | 99 | 100 | 100 | 80 | 0 | 0 | 0 | 0 |
| air | 1 | Donor | IN | 21 | 26 | 26 | BDL | 33 | 81 | 92 | 98 | | 81 | 18 | 7 | 0 | | |
| air | 1 | Naïve Control | IN | BDL | BDL | BDL | BDL | 27 | | | | | 100 | | | | | 0 |
| air | 1 | Sentinel 1 | IN | BDL | BDL | BDL | BDL | BDL | | | | | | | | | | |
| contact | 1 | Donor | IN | 27 | 26 | 35 | BDL | BDL | 98 | 95 | 98 | | | 0 | 0 | 0 | | |

| | | | | | | | | | | | | | | | | | | |
|---|---|---|---|---|---|---|---|---|---|---|---|---|---|---|---|---|---|---|
| contact | 1 | Naïve Control | IN | 32 | BDL | BDL | BDL | BDL | 100 | | | | | 0 | | | | |
| contact | 1 | Sentinel 1 | IN | 28 | BDL | BDL | BDL | BDL | 100 | | | | | 0 | | | | |
| air | 2 | Donor | IN | 26 | 25 | BDL | BDL | BDL | 84 | 93 | | | | 15 | 0 | | | |
| air | 2 | Naïve Control | IN | BDL | BDL | BDL | BDL | 33 | | | | | 100 | | | | | 0 |
| air | 2 | Sentinel 1 | IN | BDL | BDL | BDL | BDL | BDL | | | | | | | | | | |
| contact | 2 | Donor | IN | BDL | 35 | BDL | BDL | BDL | | 76 | | | | | 23 | | | |
| contact | 2 | Naïve Control | IN | BDL | BDL | BDL | BDL | BDL | | | | | | | | | | |
| contact | 2 | Sentinel 1 | IN | BDL | 36 | BDL | BDL | 27 | | 100 | | | | | 0 | | | |
| air | 3 | Donor | IN | 28 | 29 | BDL | 31 | BDL | 63 | 84 | | | | 36 | 15 | | | |
| air | 3 | Naïve Control | IN | BDL | BDL | BDL | BDL | BDL | | | | | | | | | | |
| air | 3 | Sentinel 1 | IN | 24 | BDL | BDL | BDL | BDL | 75 | | | | | 24 | | | | |
| contact | 3 | Donor | IN | BDL | 31 | 32 | BDL | BDL | | 99 | 100 | | | | 0 | 0 | | |
| contact | 3 | Naïve Control | IN | 28 | BDL | BDL | BDL | BDL | 97 | | | | | 0 | | | | |
| contact | 3 | Sentinel 1 | IN | BDL | BDL | BDL | BDL | 32 | | | | | 100 | | | | | 0 |
| air | 1 | Donor | Naïve | 25 | 26 | 27 | 14 | 23 | 83 | 92 | 99 | 100 | 99 | 16 | 7 | 0 | 0 | 0 |
| air | 1 | Sentinel 1 | Naïve | 24 | 26 | 24 | 20 | 21 | 100 | 90 | 84 | 100 | 99 | 0 | 9 | 15 | 0 | 0 |
| air | 1 | Sentinel 2 | Naïve | 32 | 27 | 24 | BDL | 29 | 100 | 99 | 100 | | 100 | 0 | 0 | 0 | | 0 |
| air | 1 | Sentinel 3 | Naïve | BDL | BDL | 33 | BDL | BDL | | | 100 | | | | | 0 | | |
| contact | 1 | Donor | Naïve | 22 | 25 | 27 | 25 | 33 | 93 | 98 | 98 | 100 | 97 | 0 | 0 | 0 | 0 | 0 |
| contact | 1 | Sentinel 1 | Naïve | 25 | 25 | 26 | 18 | 16 | 96 | 99 | 99 | 99 | 100 | 0 | 0 | 0 | 0 | 0 |
| contact | 1 | Sentinel 2 | Naïve | 33 | 25 | 25 | 23 | 21 | 83 | 100 | 99 | | 100 | 16 | 0 | 0 | | 0 |
| contact | 1 | Sentinel 3 | Naïve | BDL | 26 | 26 | BDL | 32 | | 100 | 100 | | 100 | | 0 | 0 | | 0 |
| air | 2 | Donor | Naïve | 23 | 23 | 30 | 16 | 29 | 82 | 85 | 99 | 99 | 98 | 17 | 14 | 0 | 0 | 0 |
| air | 2 | Sentinel 1 | Naïve | 25 | 22 | 27 | 19 | 22 | 87 | 100 | 100 | 99 | 100 | 12 | 0 | 0 | 0 | 0 |
| air | 2 | Sentinel 2 | Naïve | BDL | 34 | 24 | 20 | 23 | | 100 | 100 | 100 | 100 | | 0 | 0 | 0 | 0 |
| air | 2 | Sentinel 3 | Naïve | BDL | BDL | 24 | 24 | 22 | | | 99 | | 100 | | 0 | 0 | | 0 |
| contact | 2 | Donor | Naïve | 34 | 26 | 28 | 32 | 27 | 100 | 97 | 97 | 100 | 99 | 0 | 0 | 0 | 0 | 0 |

| | | | | | | | | | | | | | | | | | | |
|---|---|---|---|---|---|---|---|---|---|---|---|---|---|---|---|---|---|---|
| contact | 2 | Sentinel 1 | Naïve | 29 | 25 | 24 | 15 | 24 | 100 | 99 | 99 | 100 | 99 | 0 | 0 | 0 | 0 | 0 |
| contact | 2 | Sentinel 2 | Naïve | 28 | 30 | 27 | 17 | 23 | 100 | 100 | 99 | 100 | 100 | 0 | 0 | 0 | 0 | 0 |
| air | 3 | Donor | Naïve | 36 | 25 | 26 | 19 | 20 | 100 | 99 | 99 | 100 | 99 | 0 | 0 | 0 | 0 | 0 |
| air | 3 | Sentinel 1 | Naïve | 33 | BDL | BDL | BDL | BDL | 100 | | | | | 0 | | | | |
| air | 3 | Sentinel 2 | Naïve | BDL | BDL | BDL | BDL | BDL | | | | | | | | | | |
| contact | 3 | Donor | Naïve | BDL | 25 | 28 | 21 | 16 | | 100 | 99 | 100 | 100 | | 0 | 0 | 0 | 0 |
| contact | 3 | Sentinel 1 | Naïve | BDL | 27 | 27 | 17 | 17 | | 99 | 99 | 99 | | | 0 | 0 | 0 | |
| contact | 3 | Sentinel 2 | Naïve | BDL | 24 | 30 | 18 | 25 | | 97 | 99 | 100 | 100 | | 0 | 0 | 0 | 0 |
| contact | 3 | Sentinel 3 | Naïve | BDL | 28 | 25 | 20 | 27 | | 100 | 99 | 100 | 100 | | 0 | 0 | 0 | 0 |
| air | 1 | Donor | PI | 32 | 35 | BDL | BDL | BDL | 0 | 0 | | | | 93 | 95 | | | |
| air | 1 | Naïve Control | PI | BDL | BDL | 32 | BDL | BDL | | | 100 | | | | | 0 | | |
| air | 1 | Sentinel 1 | PI | BDL | BDL | BDL | BDL | BDL | | | | | | | | | | |
| contact | 1 | Donor | PI | 37 | 35 | BDL | BDL | BDL | | 100 | | | | 0 | 0 | | | |
| contact | 1 | Naïve Control | PI | 29 | BDL | BDL | BDL | BDL | | | | | | | | | | |
| contact | 1 | Sentinel 1 | PI | 33 | BDL | BDL | BDL | 39 | 86 | | | | | 13 | | | | 0 |
| air | 2 | Donor | PI | 29 | BDL | BDL | BDL | 37 | 0 | | | | | 95 | 99 | | | 0 |
| air | 2 | Naïve Control | PI | BDL | BDL | BDL | BDL | BDL | | | | | | | | | | |
| air | 2 | Sentinel 1 | PI | BDL | BDL | BDL | BDL | BDL | | | | | | | | | | |
| contact | 2 | Donor | PI | 23 | 30 | 37 | BDL | 35 | 95 | 96 | 57 | | 95 | 0 | 0 | 42 | | 0 |
| contact | 2 | Naïve Control | PI | BDL | BDL | BDL | BDL | BDL | | | | | | | | | | |
| contact | 2 | Sentinel 1 | PI | BDL | BDL | BDL | BDL | BDL | | | | | | | | | | |
| air | 3 | Donor | PI | BDL | BDL | BDL | BDL | BDL | | | | | | | | | | |
| air | 3 | Naïve Control | PI | BDL | BDL | BDL | BDL | BDL | | | | | | | | | | |
| air | 3 | Sentinel 1 | PI | 32 | BDL | BDL | BDL | BDL | 91 | | | | | 8 | | | | |
| contact | 3 | Donor | PI | 25 | BDL | BDL | BDL | BDL | 100 | | | | | 0 | | | | |
| contact | 3 | Naïve Control | PI | 30 | BDL | BDL | BDL | BDL | 100 | | | | | 0 | | | | |
| contact | 3 | Sentinel 1 | PI | 25 | BDL | BDL | BDL | 32 | 100 | | | | | 0 | | | | |

**Supplemental Table 3:** SARS-CoV-2 isolates used in this study.

| Virus | WHO Description | PANGO Lineage | GiSAID/Genbank Acc |
|---|---|---|---|
| SARS-CoV-2/human/USA/WA-CDC-WA1/2020 | Lineage A | WA1 | MN985325 |
| England/204820464/2020 | Alpha | B.1.1.7 | EPI_ISL_683466 |
| USA/MD-HP01542/2021 | Beta | B.1.351 | EPI_ISL_890360 |
| hCoV-19/USA/MD-HP03867/2021 | Gamma | P.1 | EPI_ISL_1468644 |
| hCoV-19/USA/KY-CDC-2-4242084/2021 | Delta | B.1.617.2 | EPI_ISL_1823618 |
| hCoV-19/USA/GA-EHC-2811C/2021 | Omicron | B.1.1.529 | EPI_ISL_7171744 |

**Supplemental Table 4:** Pathological Assessment of IN and IT inoculated Syrian hamsters on day 5. nsf = no significant findings. y = yes. n = no.

| Group | IN | | | | | | IT | | | | | |
|---|---|---|---|---|---|---|---|---|---|---|---|---|
| lesions visible grossly | n | n | n | y | n | n | y | y | y | y | y | y |
| Percentage of total area affected | | <5 | | 10 | 5 | | 50 | 90 | 70 | 50 | 70 | 60 |
| Alveolar inflammation or exudate | n | n | 1 | 2 | 1 | 1 | 4 | 4 | 4 | 4 | 4 | 4 |
| Bronchiolar epithelial cell inflammation/necrosis | n | 1 | 1 | n | n | n | 1 | n | n | n | n | n |
| Endotheleitis, vasculitis | n | n | n | y | n | n | y | y | y | y | y | y |
| Hemorrhage, fibrin and or edema | n | n | n | n | n | n | n | 3 | 3 | n | 2 | n |
| SARS2 IHC bronchioles | 0 | 1 | 1 | 1 | 1 | 1 | 0 | 1 | 0 | 0 | 0 | 0 |
| SARS2 IHC alveoli | 0 | 1 | 1 | 2 | 1 | 1 | 4 | 4 | 4 | 4 | 4 | 4 |
